# GGA2-depletion in beta cells by Enteroviruses causes Golgi acidification, premature activation of cathepsins and alters the MHC class I immunopeptidome

**DOI:** 10.1101/2025.03.28.645506

**Authors:** Klaus-Peter Knoch, Antonia Kosok, Zuzana Marinicova, Christiana Lekka, Inna Kalaidzidis, Yannis Kalaidzidis, Marko Barovic, Antje Petzold, Anke Sönmez, Carolin Wegbrod, Katharina Ganss, Andreas Müller, Alexia Carre, Raphael Scharfmann, Roberto Mallone, Michael Ghosh, Juliane Merl-Pham, Stefanie M. Hauck, Noel G. Morgan, Sarah J. Richardson, Michele Solimena

## Abstract

Enteroviruses (EVs), such as Coxsackievirus B5 (CVB5), are linked to pancreatic beta cell autoimmunity in type 1 diabetes (T1D). CVB5 infection of beta cells blocks their cap-dependent translation but spares cap-independent translation of insulin secretory granule (SG) cargoes, including major T1D autoantigens. Despite this, insulin SG stores are depleted. We show that CVB5 protease 2A rapidly depletes Golgi-associated sorting factor GGA2 in beta cells due to its short half-life. Immunostaining of pancreas sections from recent-onset T1D donors confirmed reduced GGA2 in beta cells expressing EV capsid protein VP1. Both GGA2 depletion and CVB5 2A expression impair SG biogenesis at the TGN without affecting MHC class I trafficking. They also disrupt the sorting of vacuolar ATPase and cathepsins, leading to TGN acidification and premature lysosomal hydrolase activation, which typically generate peptides presented in HLA-II alleles. Notably, the unique immunopeptidome of EV-infected, GGA2-depleted ECN90 beta-like cells is mainly presented via HLA-B alleles with an average isoelectric point (pI) of ∼5, compared to a pI of ∼7 of the prevailing HLA-I A allele immunopeptidome of non-infected cells. We propose that delayed sorting at the acidified TGN promotes cathepsin-mediated processing of in-transit secretory proteins, including SG cargoes, thereby generating novel peptides that may replace HLA class I antigens due to the acidic environment. This novel cross-presentation mechanism could have implications for T1D pathogenesis and antigen presentation in general.

## Introduction

Type 1 diabetes (T1D) results from autoimmunity directed against the insulin-producing beta cells of the pancreatic islets. While its precise cause remains unknown, T1D is thought to be the consequence of an unfavourable combination of genetic predisposition and environmental agents, such as Enteroviruses (EVs). The EV genus includes the subgroup of Coxsackieviruses (CV) and belong to the large family of picornaviruses, which are all single-stranded plus-sense RNA viruses with a genome of ∼ 7,5 kb (Rueckert and Wimmer, 1984). EVs infect primarily the gastrointestinal tract, but vary in their secondary tissue tropism, with Coxsackievirus B (CVB) preferentially targeting the heart and the pancreas (Richardson et al. 2009). Their infection can be completely asymptomatic or lead to several illnesses, ranging from mild respiratory symptoms to more severe conditions like myocarditis and meningitis. An increasing body of evidence points to EVs as potential triggers of T1D (Dahl-Jørgensen, 2024). Recent studies, in particular, have shown that EV-infection changes the HLA-class I antigen repertoire presented by beta cells, conceivably exposing them to autoimmune destruction by T-cells (Vecchio et al. 2025). Yet, the molecular mechanisms by which EVs affect the biology of beta cells, possibly fostering the generation of neoantigens, remain largely unknown.

We previously showed that acute infection of mouse insulinoma MIN6 cells with CVB5 halts the cap-dependent expression of host proteins, but not the cap-independent translation of its RNA genome as well as that of preproinsulin and other precursors of secretory granule (SG) cargoes (Knoch et al, 2014). Several of these, like insulin, ICA512/IA-2, Phogrin/IA-2beta and Chromogranin A, are major targets of humoral and/or T-cell autoimmunity in T1D. Nonetheless, CVB5 infection completely empties the SG stores due to intracellular protein degradation, as insulin secretion was abolished in the absence of apoptosis.

In this study we show that the viral protease CVB5 2A, which cleaves the viral polyprotein, but also host factors such as eIF4G, is responsible for these phenotypic alterations. Specifically, we demonstrate that CVB5 2A-induced block of cap-dependent translation reduces the levels of short half-life factors involved in membrane traffic, including the ADP-ribosylation factor-binding protein GGA2, which regulates the traffic of proteins between the trans-Golgi network (TGN) and the lysosomes. Accordingly, GGA2 depletion results in the retention of vATPase and acidification of the Golgi complex lumen. This, in turn, promotes the premature activation of lysosomal cathepsins and their cleavage of in-transit SG cargo precursors. The resulting peptides may then competitively displace weak-binders to MHC class I, including viral peptides, due to the lower luminal pH. This cascade can therefore inhibit SG biogenesis and skew the MHC class I antigenic repertoire toward the presentation of neoantigens originating from SG cargoes, and thus favour beta cell autoimmunity, while facilitating the evasion of the virus from immune recognition.

## Results

### CVB5 2A^pro^ depletes the stores of insulin SG cargoes

We previously showed that Coxsackievirus B (CVB5) depletes the insulin SG stores of infected mouse insulinoma MIN6 cells and pancreatic beta cells (Knoch et al. 2014) Therefore, we searched for factor(s) responsible for this alteration, focusing on CVB5 proteases. The ∼ 7.5 Kb virus genome encodes a single viral polypeptide, which is cleaved into several proteins by CVB5 proteases 2A (2A^pro^) and 3C (3C^pro^) (Svitkin et al, 1979; Toyoda et al, 1986) (Fig. 1a). CVB5 3C^pro^ has a nuclear localisation sequence and cleaves factors associated with cellular DNA-dependent RNA polymerase I, II and III (Sharma et al, 2004; Clark et al, 1993; Yalamanchili et al, 1997), thereby impairing transcription of host cells. CVB 2A^pro^, in turn, cleaves elF4GI and eIF4GII to silence cap-dependent translation (Sommergruber et al, 1994; Gradi et al. 1998). In addition, both proteases inhibit translation by cleaving poly(A)-binding protein (Pabp) (Kuyumcu-Martinez et al, 2002). We first verified that overexpressed CVB5 2A^pro^ and 3C^pro^ in mouse insulinoma MIN6 cells cleave Pabp and eIf4g (Suppl. Fig. 1a). Notably, only CVB5 2A^pro^ overexpression reduced the levels of insulin SG markers Ica512, Pc1/3 and Pc2, mimicking the phenotype observed upon CVB5 infection (Knoch et al, 2014) (Fig. 1b). Mass spectrometric analyses validated that in CVB5 2A^pro+^ cell extracts the levels of SG cargoes were reduced (Fig. 1c). Comparable alterations were observed in cells overexpressing 2A^pro^ of Echovirus-9 DM (EV-9 DM) - an enterovirus isolated from a subject deceased shortly after the onset of T1D (Paananen et al, 2003) (Suppl. Fig. 1b). We further measured the levels of proinsulin and insulin in MIN6 cell extracts by ELISA. Neither protease prevented the glucose-stimulated upregulation of proinsulin (Fig. 1d), consistent with its reliance on cap-independent translation (Knoch et al, 2014). In contrast, in CVB5 2A^pro+^ cells the levels of mature insulin were 60% lower compared to control cells (Fig. 1e). Accordingly, the detection of SGs by insulin immunostaining in cells double positive for CVB5 2A^pro^ and GFP as a reporter for transfection was strongly reduced relative to the adjacent CVB5 2A^pro-^/GFP^-^ cells (Fig.1f), although labelling persisted in the perinuclear region, compatible with the detection of proinsulin in-transit through the Golgi complex (Fig. 1f). Being a secretory protein insulin is unlikely to be the target of proteasomal degradation and all three proteasomal pathways, namely trypsin-like, chymotrypsin-like and caspase-like activities were not significantly increased in CVB5 2A^pro+^ cells (Suppl. Fig. 1c). Likewise, proteasome inhibition with MG132 (Suppl. Fig. 1d) did not prevent the CVB5 2A^pro^-induced loss of mature SG proteins. Moreover, CVB5 2A^pro+^ cells do not express the subunits Psmb8, 9 and 10 of the immunoproteasome (Suppl. Fig. 1e). We further tested the occurrence of apoptosis. Neither CVB5 2A^pro^ nor 3C^pro^ overexpression activated Caspase3 and Caspase7 (Suppl. Fig. 1f), nor induced Parp cleavage (Suppl. Fig. 1g). Taken together, these data indicate that the degradation of mature SG cargoes in CVB5 2A^pro+^ cells does not involve the proteasome and must occur along the secretory pathway.

**Figure 1:**
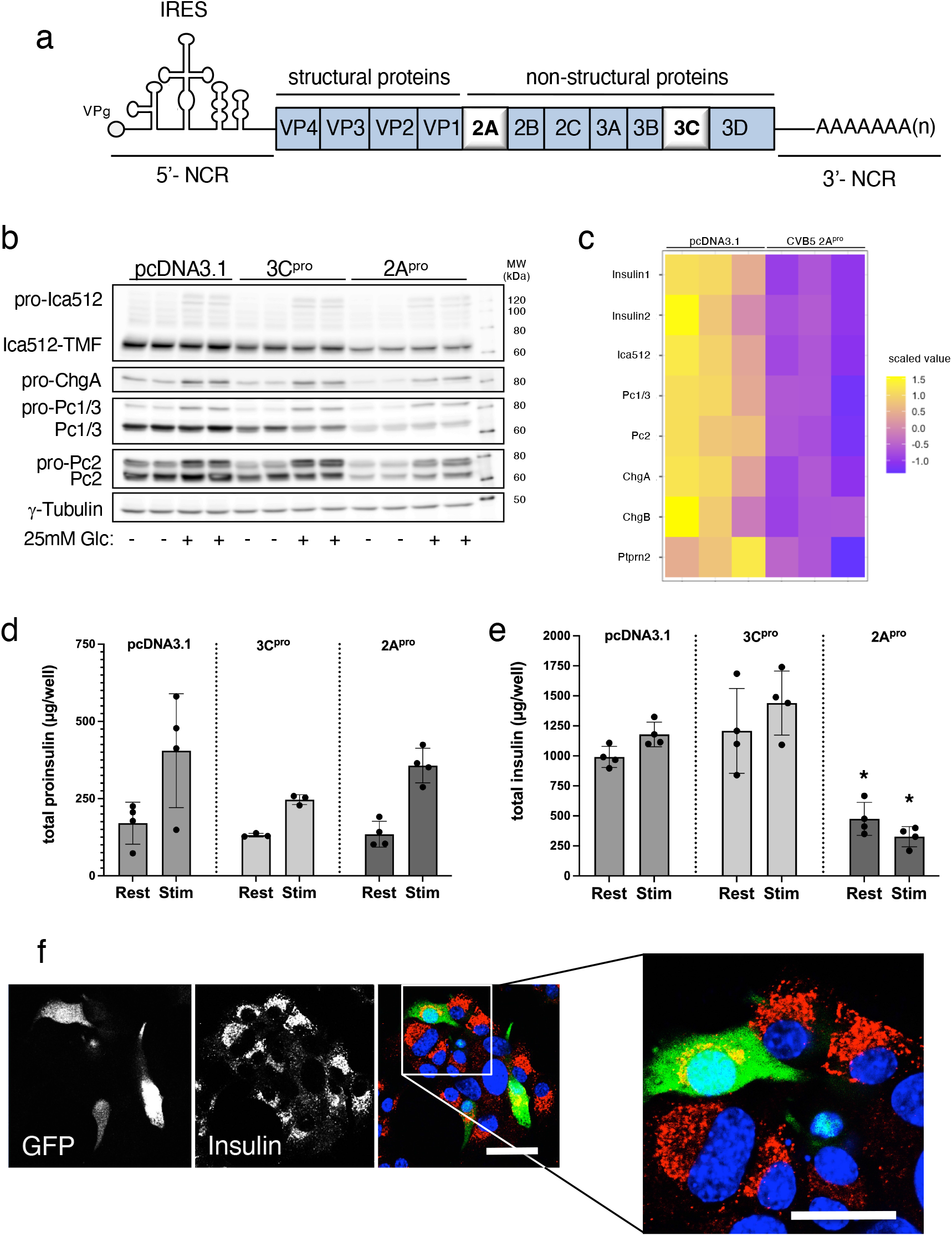
CVB5 2A, but not 3C, depletes insulin granule stores. (a) Scheme of the picornavirus genome. IRES: internal-ribosome entry site. (b) Western blot for Ica512, Chga, Pc1/3, Pc2 and ɣ-Tubulin in extracts of MIN6 cells transfected with either CVB5 2A or 3C proteases or the empty vector pcDNA3.1. (c) Heatmap for insulin1, insulin2, Ica512, Pc1/3, Pc2, Chga and Chgb and Ptpn2 from global proteomic analysis (Z-score clustered) of extracts of MIN6 cells transfected with CVB5 2A or pcDNA3.1 alone. (d, e) Quantification by ELISA of proinsulin (d) and insulin (e) in the media and extracts of MIN6 cells transfected with either CVB5 2A or 3C. Data are from ≥ 3 independent experiments. *p<0.05 by one-way ANOVA. (f) Immunostaining of insulin in MIN6 cells co-transfected with CVB5 2A and GFP as a transfection reporter. Nuclei are counterstained with DAPI. Scale bars = 20 µm.

### CVB5 2A^pro^ targets insulin SG cargoes to degradation along the secretory pathway

To test this hypothesis further we analyzed by radioactive pulse-chase labelling the half-life of proIca512 and proPc1/3 in control and CVB5 2A^pro+^ MIN6 cells. Conversion of these proforms following their exit from the Golgi complex and sorting into immature SGs should result in the decrease of their levels, with an equivalent increase of the corresponding mature forms. In control cells and CVB5 2A^pro+^ cells the half-life of proIca512 and proPc1/3 were comparable (Fig. 2a and 1b). However, in CVB5 2A^pro+^ cells the appearance of mature Ica512, termed Ica512-TMF, and Pc1/3 was significantly delayed compared to control cells (Fig. 2c and 1d). We also exploited the pulse-chase labelling of Insulin-SNAP with TMR to detect newly-synthesized SGs (Ivanova et al, 2013; Kempter et al, 2021) (Fig. 2e). Lack of Insulin-SNAP^TMR^ in CVB5 2A^pro+^ MIN6 cells corroborated the conclusion that the protease alters the traffic of SG cargoes, leading to their degradation at the time when SGs originate from the trans-Golgi network (TGN).

**Figure 2:**
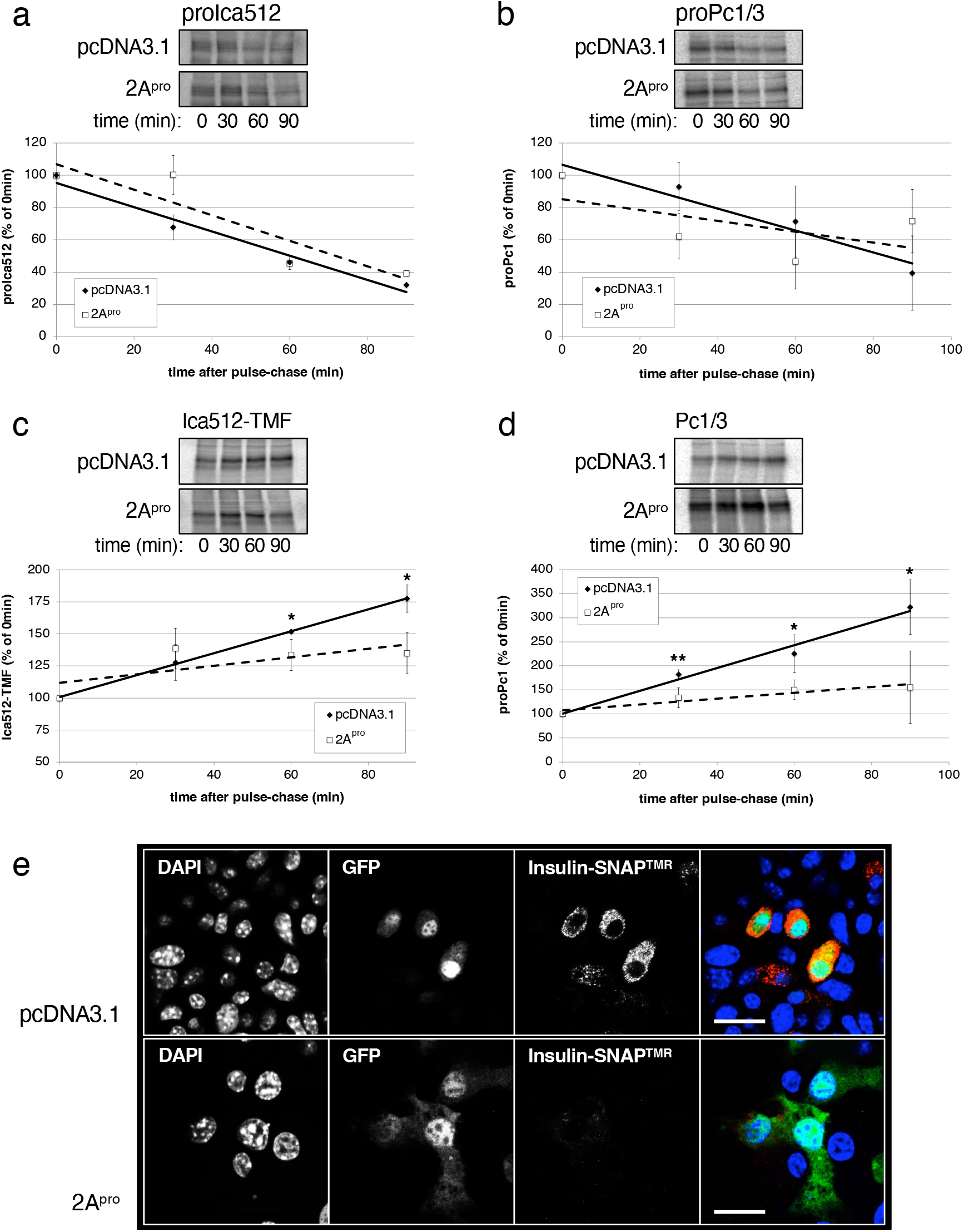
CVB5 2A delays the maturation of insulin granule proteins. (a-d) Radioactive pulse-chase labeling of Ica512 (a and c) and Pc1/3 (b and d) in MIN6 cells 30, 60 and 120 min after stimulation with 25 mM glucose following 1 hour resting with 0 mM glucose. Data are from 3 independent experiments. (e) TMR labeling (in red) of newly synthesized insulin in MIN6 cells co-transfected with CVB5 2A, Insulin-SNAP and GFP as transfection reporter. Nuclei are counterstained with DAPI. Scale bar = 20 µm.

### CVB5 2A^pro^ reduces the levels of Gga2 and Rab3b

To identify possible factors involved in the CVB5 2A^pro^-induced depletion of SG stores, we screened the mass spectrometry proteomic data from control, CVB5 2A^pro+^ and EV-9 DM 2A^pro+^ MIN6 cells for changes in the levels of cytosolic proteins involved in post-Golgi trafficking. Nine of such proteins in our list were reduced by overexpression of either CVB5 2A^pro^ (Fig. 3a) or EV-9 DM 2A^pro^ (Suppl. Fig. 2a). Analysis by immunoblotting of CVB5 2A^pro+^ MIN6 cell extracts validated the reduction of four of these proteins, namely Arf1, Sytl4, Rab3b and Gga2 (Fig. 3b-c), while in the case of EV-9 DM 2A^pro+^ MIN6 cell extracts we observed a lower content for Gga2, Rab3b and Vtl1b (Suppl. Fig 2b-c). Quantitative RT-PCR assays indicated that CVB5 2A^pro^ had no impact on the expression levels of *Arf1, Sytl4, Rab3b* and *Gga2* mRNAs (Fig. 3d). Next, we analyzed the half-life of these four proteins by blocking their translation with cycloheximide. Notably, the levels of Rab3b and Gga2, but not those of Arf1 and Sytl4, were reduced by ∼75% 24 hours after cycloheximide treatment (Fig. 3e-f), indicating that both Rab3b and Gga2 have a short half-life. This property may account for their rapid depletion upon CVB5 2A^pro^-induced block of cap-dependent translation. Such conclusion was corroborated by evidence that neither purified Rab3b-GST nor Gga2-GST levels were affected upon incubation with CVB5 2A^pro+^ MIN6 cell extracts for up to 6 hours (Suppl. Fig. 2d), while eIf4g was cleaved, suggesting that Rab3b and Gga2 are not proteolytic substrates of CVB5 2A^pro^. Since Gga2 and Rab3b were the only two post-Golgi trafficking proteins in our list depleted by both CVB5 2A^pro^ and EV-9 DM 2A^pro^ and each having a short half-life, the following studies were restricted to analyze their relationship with the CVB5 2A^pro^-induced exhaustion of the SG stores.

**Figure 3:**
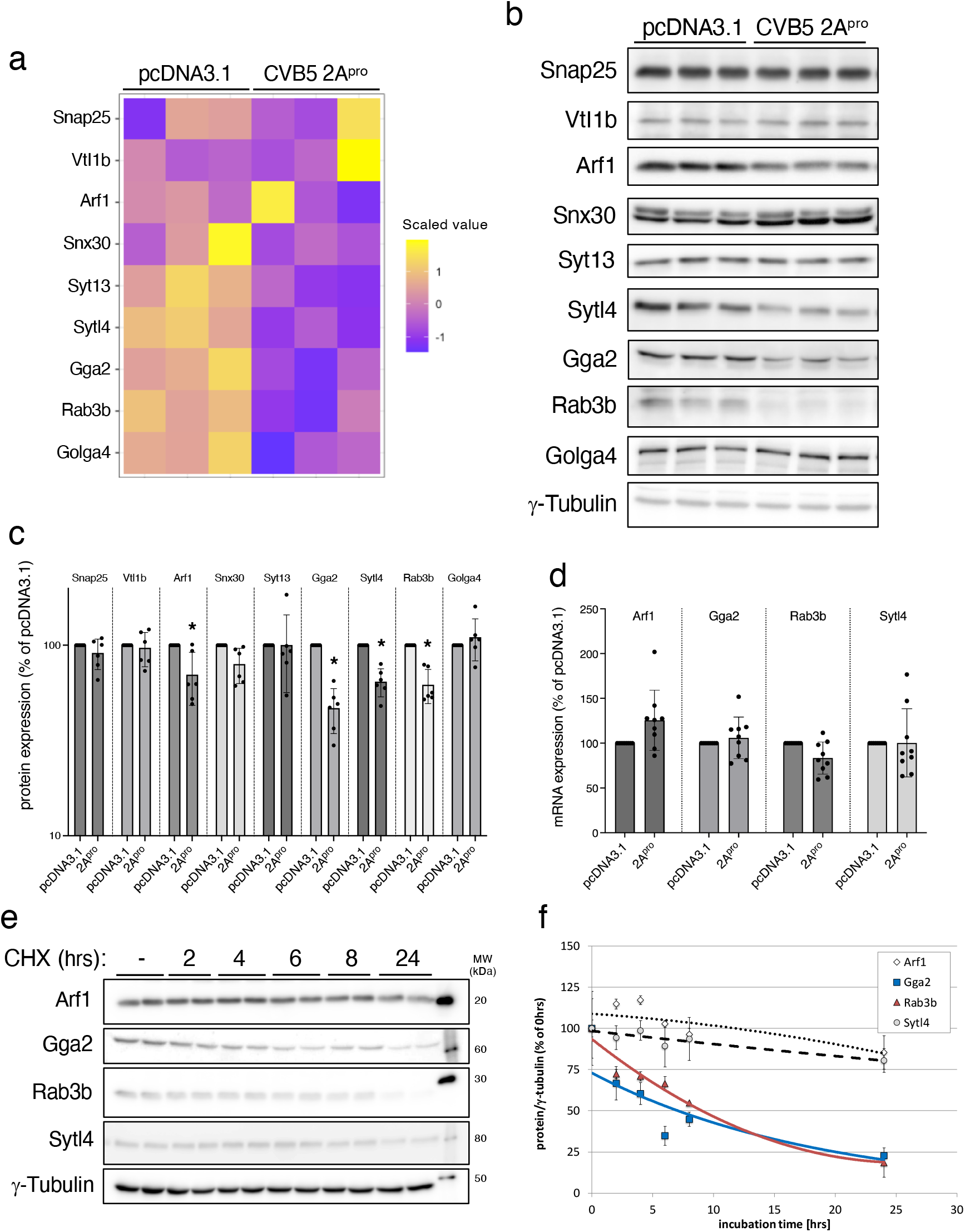
CVB5 2A depletes multiple components of the post-Golgi trafficking machinery. (a) Heatmap for selected players in post-Golgi membrane trafficking as revealed by global proteomic analysis (Z-score clustered) of MIN6 cells transfected with CVB 2A or pcDNA3.1 alone. (b, c) Western blots (b) and corresponding quantifications (c) for selected post-Golgi trafficking proteins downregulated upon expression of CVB5 2A. (d) Quantitative RT-PCR for *Arf1, Gga2, Rab3b* and *Sytl4* in MIN6 cells transfected with CVB 2A or pcDNA3.1 alone. mRNA expression levels were normalised by β-actin mRNA. Values are shown as percentage of control cells. (e,f) Western blots (e) and corresponding quantifications of the half-life (f) of the indicated proteins in MIN6 cells treated with cycloheximide (100µg/ml) to block protein biosynthesis. All data are from ≥ 3 independent experiments; statistical values were calculated by one-way ANOVA *p<0.05.

### Silencing of Gga2 depletes SG stores similarly to CVB5 2A^pro^

Gga2 is a member of the Golgi-localized, gamma adaptin ear-containing, ARF-binding protein family which binds to the mannose-6-phosphate receptor and is required for the sorting at the Golgi complex of lysosomal hydrolases, including cathepsins, for their delivery to the endo/lysosomal compartment (Zhu et al. 2001, Hida et al. 2007). The small GTPase Rab3b, in turn, plays a role in membrane trafficking and regulated exocytosis of secretory vesicles, being especially enriched on newly-synthesised SGs (Neukam et al, 2024). To investigate whether depletion of either Gga2 or Rab3b, similarly to CVB5 2A^pro^, reduced SGs stores, we silenced their expression in MIN6 cells. The effectiveness of these knockdowns was validated by qPCR (Fig. 4a) The knockdown was specific for the proteins and the cell did not compensate for the loss of Gga2 nor Rab3b with up-regulation of one of the other family members (Suppl. Fig. 3a, Suppl. Fig. 3b). Upon *Gga2* knockdown, the levels of Rab3b were reduced (Fig. 4b). Conversely, *Rab3b* knockdown did not alter the levels of Gga2. Neither knockdowns affected the levels of pro-Ica512, and proPc1/3 (Fig. 4b) in either glucose-resting or -stimulated conditions compared to control cells, while those of proPc2 were reduced regardless of the glucose levels. On the contrary, mature Ica512/Ica512-TMF, Pc1/3 and Pc2 were only reduced upon knockdown of *Gga2* (Fig. 4b). Total proinsulin levels were reduced in resting and glucose-stimulated *Gga2* siRNA-treated cells, but only in glucose-stimulated *Rab3* siRNA-treated cells (Fig. 4c). A trend for reduction of insulin was observed upon silencing of *Gga2* or *Rab3* (Fig. 4d). As expected, glucose-stimulated insulin secretion was impaired upon *Rab3b* depletion, while only a trend for reduction was observed upon silencing of *Gga2* (Fig. 4e). In both instances the immunoreactivity for insulin in the perinuclear region was enhanced, compatible with proinsulin accumulation in the Golgi complex, while the peripheral granular staining was reduced (Fig. 4f). Since Rab3b plays a role in post-Golgi trafficking steps that are distal to those involving Gga2 at the trans-Golgi network (TGN), and silencing of the latter more closely resembled the traits observed in CVB5 infected and CVB5 2A^pro+^ cells, with lower levels of SG cargoes, we further focused our analysis on Gga2 only.

**Figure 4:**
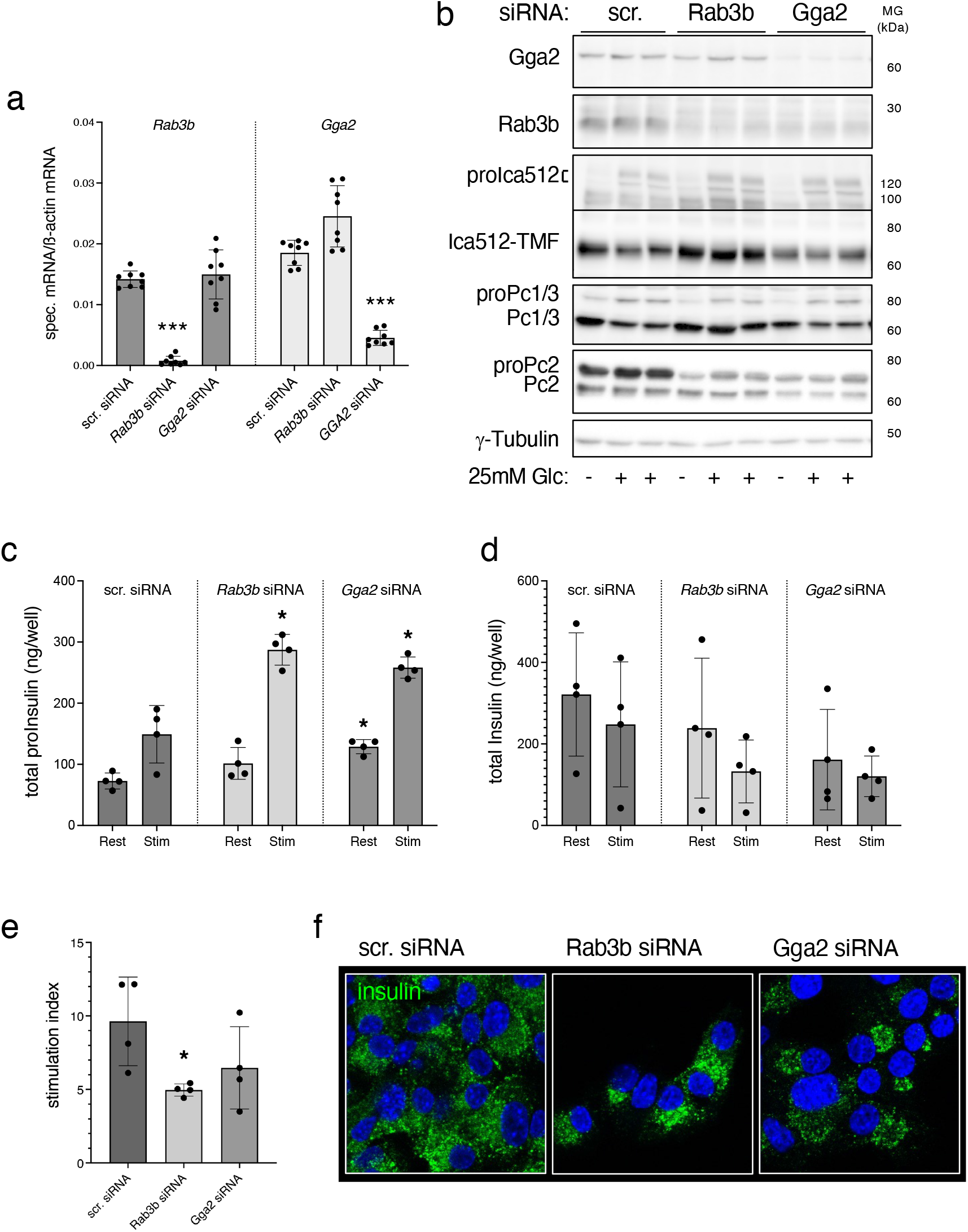
The phenotypes of *RAB3b* and *GGA2* depleted MIN6 cells resemble that of CVB5 2A-expressing MIN6 cells. (a) Quantification of *Gga2, Rab3b* and *Sytl4* mRNA levels in MIN6 cells treated with siRNAs for the corresponding genes, as measured by RT-PCR. Values were normalised to *β-Actin* mRNA levels and shown as percentage of those in control cells treated with a scrambled siRNA (n=3). (b) Western blots for Rab3b, Gga2, Sytl4, Ica512, Pc1/3, Pc2 and ɣ-Tubulin in extracts of MIN6 cells treated with *Gga2, Rab3b, Sytl4* or *scrambled* siRNAs and incubated with 0 (-) or 25 mM (+) glucose. (c,d) Quantification by ELISA of proinsulin (c) and insulin (d) amounts in the media and extracts of MIN6 cells treated as in (b). (e) Insulin stimulation index of MIN6 cells treated as in (b). Immunostaining of insulin in *Rab3b* or *Gga2* siRNA depleted MIN6 cells. Nuclei are counterstained with DAPI. Scale bar = 20 µm. All data are from ≥ 3 independent experiments; statistical values were calculated by one-way ANOVA. *p<0.05.

### GGA2 levels are reduced in EV VP1^+^ beta cells of donors with T1D

We searched for evidence of GGA2 downregulation in pancreatic beta cells of subjects deceased shortly after the onset of T1D and positive for the expression of the Coxsackievirus protein VP1. To this aim, pancreas tissue sections of these donors were immunostained with anti-VP1, anti-insulin and anti-GGA2 antibodies and the corresponding signals in confocal images were analyzed with HALO to assess the distribution of insulin and GGA2 in VP1^+^ cells (Fig. 5a and Suppl. Fig. 4a). The signals for insulin and GGA2 showed a clear direct proportionality, with the GGA2 signal being higher in insulin^+^ cells compared to Insulin^-^ cells (Suppl. Fig. 4b). In VP1^+^ cells the signal for Insulin was reduced by 70% compared to VP1^-^ cells (Fig. 5b), consistent with the notion that EV infection of beta cells correlates with their insulin content being lower. Remarkably, the signal for GGA2 in VP1^-^ beta cells was also significantly higher than in VP1^+^ cells (Fig. 5c). These data in human beta cells in situ corroborate the findings in MIN6 cells, indicating that block of cap-dependent translation by CVB5 2A^pro^ rapidly depletes beta cells of Gga2.

**Figure 5:**
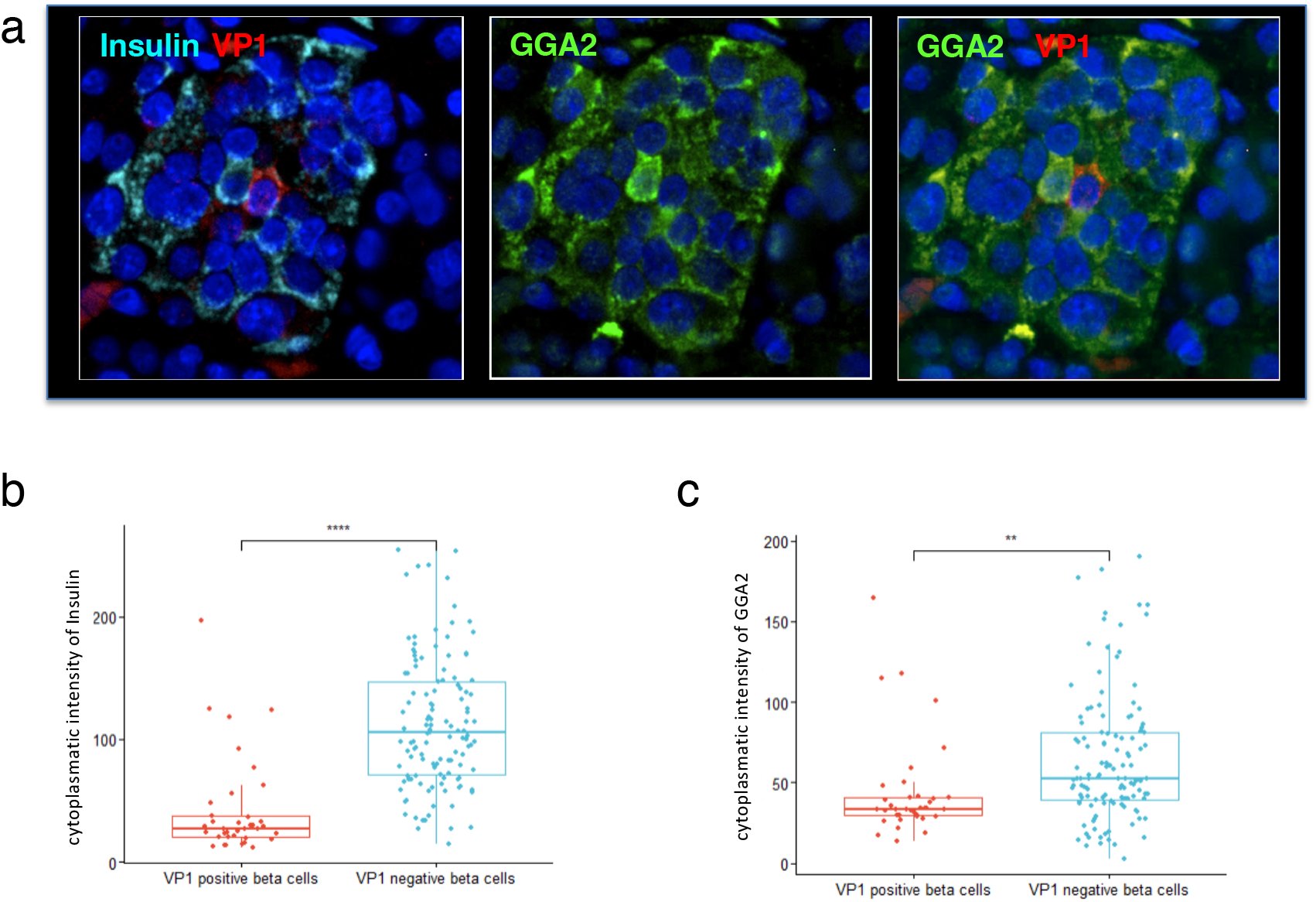
Enterovirus infection correlates with reduced insulin and GGA2 expression in human pancreatic tissue of donors with T1D. (a) Immunostaining of insulin (cyan), GGA2 (green) and VP1 (red) in human tissue sections. Confocal images were input into HALO. Using an annotation tool, uniform square-shaped annotations were created. These were randomly annotated in 8-9 areas within VP1^+^ beta cells (representative image in top panel) and 8-9 areas in VP1^-^ beta cells (representative image in bottom panel). Using the area quantification module within HALO, the average GGA2 staining intensity was calculated and presented in the box plot. Quantification of (b) insulin and (c) GGA2 in VP1^+^ cells. (d) Ratio between insulin and GGA2 in human beta cells

### Gga2 knockout alters the post-Golgi trafficking of newly-synthesized insulin

Next, we investigated if deletion of *Gga2* altered post-Golgi trafficking in MIN6 cells. Neither immunostaining for the Golgi marker Golga4 (Suppl. Fig. 5a) nor electron microscopy (Suppl. Fig. 5b) revealed gross changes in the appearance of the Golgi apparatus in *Gga2*^*-/-*^ MIN6 cells. We then analyzed the distribution of cathepsins, which depend on Gga2 for their sorting to the endolysosomal compartment (Doray et al. 2020). Specifically we focused on CtsB, CtsD and CtsL, which, based on transcriptomic data, are the most abundant cathepsins in our MIN6 cells (data not shown). CtsB-GFP, CtsD-GFP, and to a lesser extent also CtsL-GFP, were enriched in the perinuclear Golgi complex region of *Gga2*^*-/-*^ MIN6 cells compared to wild type cells (Fig. 6a), suggesting that their exit from the TGN was impaired. Next, we tested whether lack of *Gga2* also affected the sorting of insulin SG cargoes, which is unknown. To image by confocal microscopy the biogenesis of insulin SGs from the TGN, wild type and *Gga2*^*-/-*^ MIN6 cells were transfected with the Insulin-SNAP reporter (Ins-SNAP). After blocking all preexisting insulin-SNAP with a non-fluorescent SNAP substrate, newly-synthesized Ins-SNAP was pulse-labelled with the fluorescent SNAP-TMR substrate at 19°C to prevent exit from the TGN, and then chased at 37 °C. In control cells at time 0 of the chase, Insulin-SNAP^TMR^ was broadly distributed throughout the cell, conceivably being still largely found in the endoplasmic reticulum (Fig. 6b). After 30 minutes, Insulin-SNAP^TMR^ was enriched in the perinuclear Golgi region and 30 minutes later the first individual Insulin-SNAP^TMR+^ objects appeared, after 2 hours Insulin-SNAP^TMR+^ was only visible in peripheral organelles resembling SGs. In *Gga2*^*-/-*^ MIN6 cells the trafficking of newly-synthesized Insulin-SNAP^TMR^ was delayed, with its signal still being detectable in the TGN area after 2 hours. Time-lapse spinning disc confocal microscopy indicated that at 60 minutes of the chase the average diameter of Insulin-SNAP^TMR+^ vesicles in control cells was 0.33±0.01 µm (mean±SEM, 50048 objects) compared to 0.37±0.01 µm (mean±SEM; 44682 objects) in *Gga2*^*-/-*^ MIN6 cells. (Fig. 6c and Suppl. Movies 1-2). Moreover, the mean distance of all detected insulin-SNAP^TMR+^ vesicles in control cells increased overtime from 3.51±0.05 µm to 3.87±0.13 µm compared to 3.45±0.04 µm to 4.08±0.1 µm in *Gga2*^-/-^ MIN6 cells, with a plateau being reached after 53 ± 6.3 minutes in wild type cells, but only after 90 ± 6.5 minutes in *Gga2*^*-/-*^ MIN6 cells (Fig. 6d), with the larger mean distance between Insulin-SNAP^TMR+^ vesicles at the end of the chase reflecting their lower total number. In contrast, the constitutive delivery of MHC class I molecules to the surface of *Gga2*^-/-^ MIN6 cells was not altered (Fig. 6e). The levels of MHC class I molecules measured by FACS at the cell surface of control and *Gga2*^*-/-*^ MIN6 cells kept either at 37 °C, at 19°C, or first at 19 °C and then again at 37 °C were also comparable (Fig. 6f).

**Figure 6:**
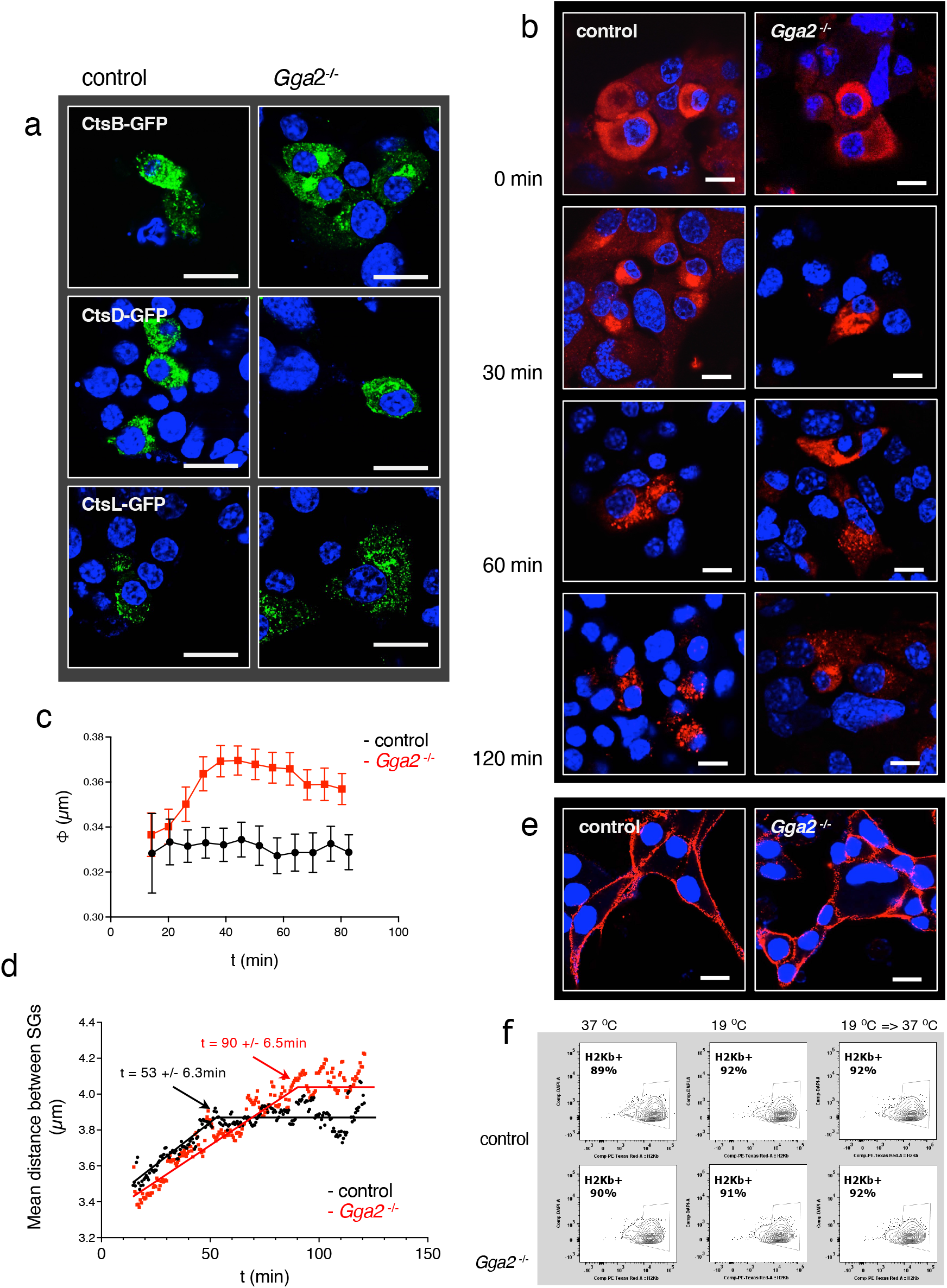
Perturbed trafficking of SG cargoes, but not of MHC-I H2Kb in *Gga2*^*-/-*^ MIN6 cells. a) Confocal microscopy imaging of newly synthesized Insulin-SNAP^TMR^ in control and *Gga2*^-/-^ MIN6 cells pulse-labeled with SNAP^TMR^ for 30’ minutes at 19 °C and then chased at 37 °C for 30, 60 or 120 minutes before fixation. Time 0 is when cells were shifted from 19 °C to 37 °C. Nuclei were counterstained with DAPI. Scale bars = 10 µm. (b, c) Quantification of the diameter (b) and mean reciprocal distance (c) of newly-generated Insulin-SNAP^TMR+^ objects in living control and *Gga2*^-/-^ MIN6 cells pulse-labeled and then chased like in (a). Data were collected from 21 and 24 time-lapse spinning disk confocal microscopy movies and analyzed with the Motion Tracking software. Time 0 is when cells were shifted from 19 °C to 37 °C. (d) Immunostaining of MHC-I H2kb in control and *Gga2*^-/-^ MIN6 cells. Nuclei were counterstained with DAPI. Scale bars = 10 µm (e) FACS analysis of H2kb immunostained control and *Gga2*^-/-^ MIN6 cells, which had been constantly kept either at 37 °C or 19 °C, or incubated at 19 °C for 90 min before been shifted to 37 °C for 120 minutes (a).

### Acidification of the Golgi complex in Gga2-depleted cells leads to premature activation of cathepsins

As depletion of *Gga2* altered the sorting of both cathepsins and SG cargoes, we wonder whether premature activation of lysosomal hydrolases in the TGN accounted for the degradation of in-transit SG cargoes. Conversion of inactive procathepsins into active cathepsins requires their self-cleavage, which normally occurs in the endolysosomes following the vATPase-driven acidification of the luminal pH <6. Notably, in *Gga2*^*-/-*^ MIN6 cells vATPase subunits Atp6v0a1-GFP and Atp6v1b2-GFP were enriched in the perinuclear region compared to control cells, suggesting that their sorting was also altered (Fig. 7a). Therefore we conjugated resident proteins of the ERGIC (lectin, mannose binding 1; Lman1/Ergic53-CFP), cis-Golgi (beta-1,4-galactosyltransferase7; Gt-GFP), late Golgi (Golgi membrane protein 1; Golm1-CFP) and TGN (trans-Golgi network integral membrane protein 1; Tgoln1-CFP) to CFP, a reliable fluorescent reporter for pH measurement in living cells by FLIM (Poëa-Guyon et al., 2013a) (Suppl. Fig. 6a). The expression and proper localization of each of the CFP-reporters in stably transfected control MIN6 cells was verified by confocal microscopy (Suppl. Fig. 6b). The linear response of the CFP-reporters was validated by equilibrating nigericin+monensin permeabilized cells in buffers with pH ranging between 5.5 and 7.5 (Suppl. Fig. 6c). Cell clones expressing the various CFP reporters were transiently co-transfected with CVB5 2A^pro^ or pcDNA3.1 together with mKate2, with the latter being a reporter for transfection. Cytosolic mKate2 did not affect the half-life of the CFP-reporters in their respective compartment along the secretory pathway (Suppl. Fig. 6d and 6e). In the clones expressing the CFP-reporters, pcDNA3.1 and mKate2 the pH of the ERGIC, cis-Golgi, late Golgi and TGN compartments corresponded to the expected mean values of 7.10 +/-0,19, 6.81 +/-0,07, 6.65 +/-0,12 and 6.21 +/-0,14, respectively. In cells cotransfected with CVB5 2A^pro^ the pH of the ERGIC was not significantly changed, whereas that of the cis-Golgi, late Golgi and TGN compartments was significantly lower than in pcDNA3.1 control cells (Fig. 7b). In *Gga2*^*-/-*^ MIN6 cells transiently transfected with the CFP reporters, the luminal pH of the late Golgi and TGN was also significantly lower than in control MIN6 cells (Fig. 7b). Next, we measured the activities of CtsB and CtsD normalized to their expression levels in Golgi enriched fractions from control and *Gga2*^*-/-*^ MIN6 cells, as well as in MIN6 cells transiently expressing pcDNA3.1^+^ or CVB5 2A^pro+^ (Fig. 7c). Both cathepsin activities were enhanced in *Gga2*^*-/-*^ and CVB5 2A^pro+^ MIN6 cells, despite CtsB and CtsD levels being reduced compared to control cells (Suppl. Fig. 6f and 6g), conceivably due to the block of their cap-dependent translation. Taken together, these data point to a scenario whereby delayed sorting of lysosomes and SG cargoes into post-Golgi vesicles upon genetic or CVB5 2A^pro^-induced depletion of Gga2 leads to the vATPase-driven acidification of the TGN, with premature activation of the retained cathepsins and degradation of in-transit, cap-independent translated SG cargoes.

**Figure 7:**
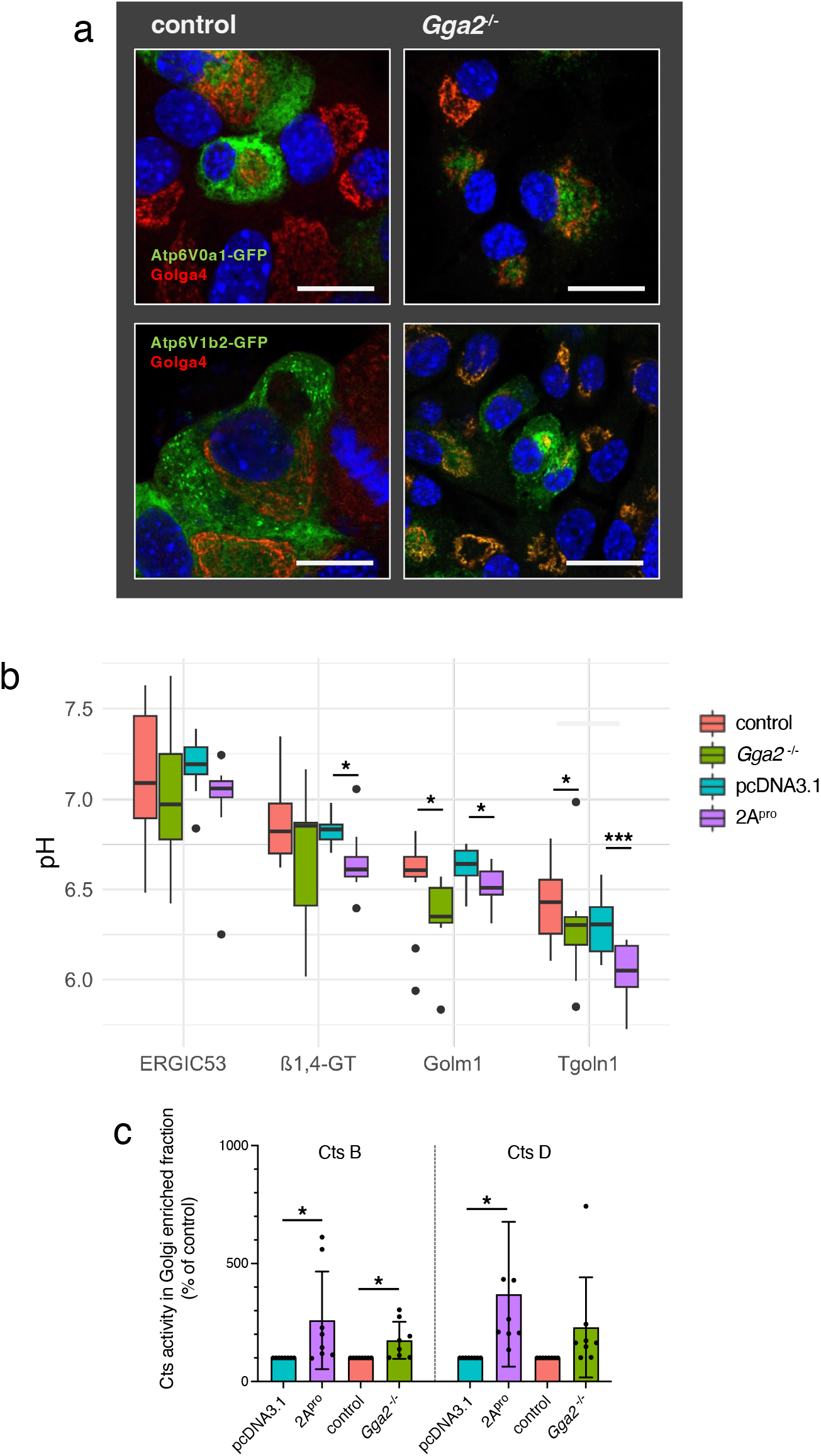
CVB5 2A and *Gga2* depletion induce the acidification of the TGN and the premature activation of Cathepsins. (a) Confocal microscopy for ATP6V0a1-GFP in wild-type and *Gga2*^-/-^ MIN6 cells. The Golgi complex was imaged by co-immunostaining for Golga4. Nuclei are counterstained with DAPI. Scale bar = 20 µm. (b,c) pH values along the early secretory pathway of wild-type MIN6 cells (b) or *Gga2*^-/-^ MIN6 cells as well as in MIN6 cells transfected either with pcDNA 3.1 or CVB 2A (c). Measurements were obtained by FLIM using markers for the ERGIC (Ergic53), cis-Golgi (1.4 beta Galactosyl Transferase, GT), late Golgi (Golm1) and TGN (Tgoln1) fused to CFP. Quantifications are from ≥8 independent experiments, statistical values were calculated by one-way ANOVA. *p<0.05. d). CtsB and CtsD activities in Golgi-enriched fractions isolated from wild-type or *Gga2*^-/-^ MIN6 cells as well as MIN6 cells transfected either with pcDNA3.1 or CVB 2A. Data are from ≥ 5 independent experiments, statistical values were calculated by one-way ANOVA. *p<0.05.

### The unique immunopeptidome of EV-9 DM infected ECN90 cells is skewed towards presentation of acid peptides in HLA-B

The scenario illustrated above could result in the generation of neoantigens to be presented by MHC class I molecules at the cell surface. To test this hypothesis we analyzed the peptidome of HLA class I molecules (HLA-I) in control and EV-9 DM-infected human beta ECN90 cells, which express the T1D susceptibility allele A*02:01. Cells were infected with EV-9 DM rather than CVB5 because the latter had been adapted for infection of mouse insulinoma cells (Al-Hello et al. 2005). Conditions of infection were optimized with regard to virus titer, number of plated cells and harvesting time in order to detect VP1 and eIF4G cleavage, while minimizing that of cleaved caspase 3, 7 and PARP1 as markers of apoptosis (Suppl. Fig. 7a, 7b). EV-9 DM infection neither changed the total levels of HLA-I as measured by qPCR (Fig. 8a) nor the protein levels of the HLA-A, HLA-B alleles and the associated β-microglobulin (Fig. 8b). It also did not alter the immunocytochemical pattern of HLA-I in VP1^+^ ECN90 cells (Fig. 8c). However, as in CVB5-infected MIN6 cells, insulin and also GGA2 were significantly reduced by 70% and 50%, respectively (Fig. 8b). Therefore, we isolated the peptidome of EV-9 DM-infected and control ECN90 cells by immunoprecipitation and subsequent LC-MS/MS. On average, the number of identified peptides eluted from HLA-I across all replicates (3 control and 3 EV-9 DM infected EC-90 cells) was 494, for a total of 3,366 peptides, including 1,113 unique peptides. The distribution of unique peptides eluted from HLA-I of EV-9 DM-infected (282) and control (337) ECN90 cells was similar (Suppl. Fig. 7c), with 56% of them (282+337) being unique for one or the other condition, and the remaining 494 (44%) being presented by both EV-9 DM-infected and control ECN90 cells (Suppl. Fig. 7c). In particular, 54/282 peptides were exclusively recovered in all 3 replicates from infected cells, whereas 13/337 peptides were recovered from all control samples only (Fig. 8d).

**Figure 8:**
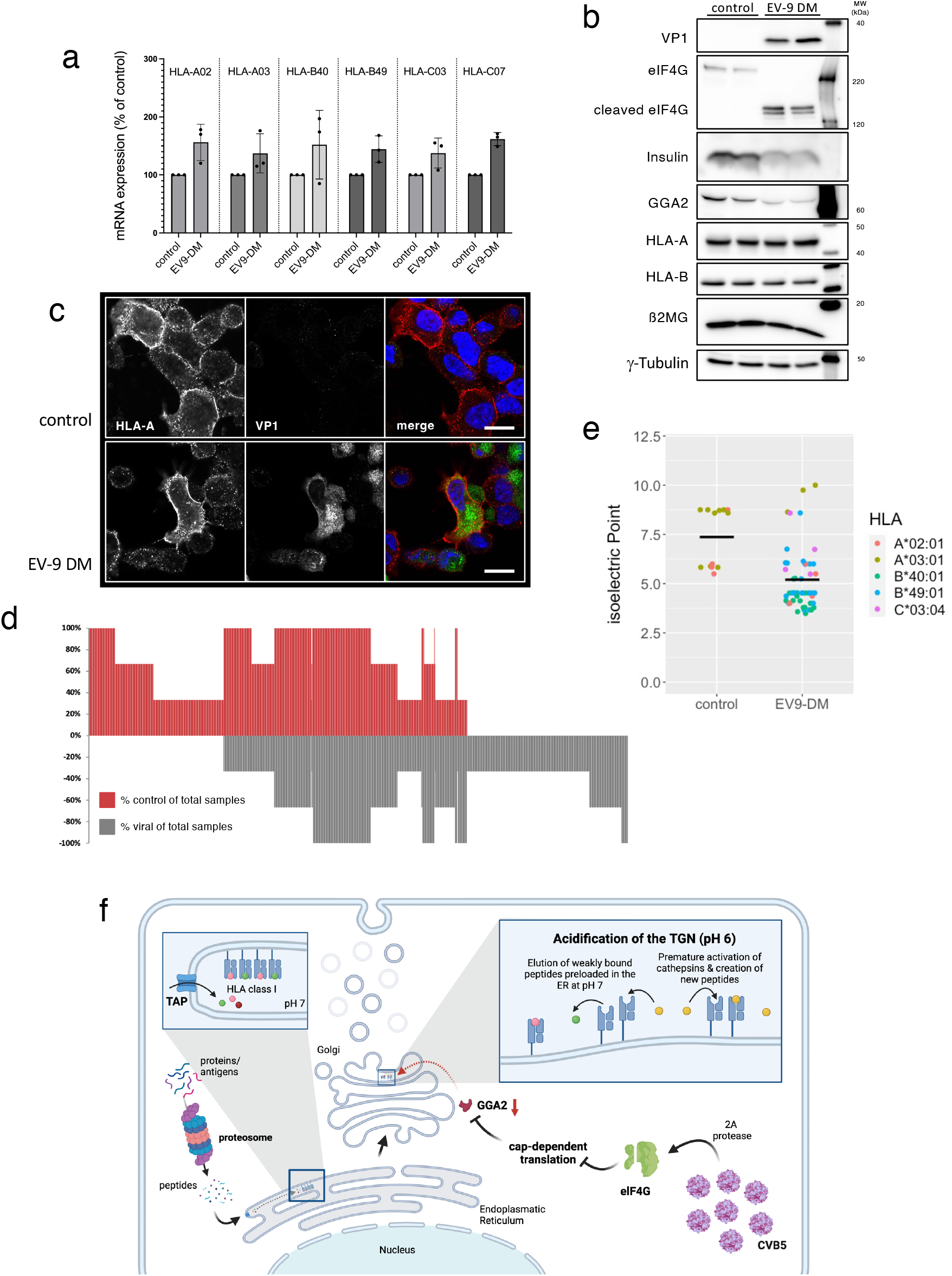
Enterovirus infection changes the peptide repertoire presented in human ECN90 insulinoma cells. ECN90 cells were infected with EV-9 DM virus and harvested 2 days after infection. (a) Quantitative RT-PCR for *HLA-A02, HLA-A03, HLA-B40, HLA-B49, HLA-C03* and *HLA-C07* mRNAs in ECN90 cells infected with EV-9 DM or uninfected as control. mRNA expression levels were normalised by b-actin mRNA. Values are shown as the percentage of uninfected control cells (n=3). (b) Western blot for VP1, eIF4G, insulin, GGA2, HLA-A, HLA-B, β2MG and ɣ-Tubulin in extracts of uninfected ECN90 cells or infected with EV-9 DM. (c) Immunostaining of HLA-B in ECN90 cells. Nuclei are counterstained with DAPI. Scale bar =10 µm. (d) Distribution of unique HLA-I peptides across all samples. Unique HLA-I-eluted peptides are arranged along the x-axis according to their presence in EV-9 DM infected ECN90 cells (red) or control samples (grey) with the peptides present exclusively in infected cells on the left-hand side and peptides exclusively in control on the right-hand side. (e) Jitter blot of the isoelectric point of the identified peptides isolated from HLA-I of ECN90 cells calculated using Compute pI/Mw software (https://web.expasy.org/compute_pi/). (f) Graphic illustration for how EV 2A^pro^ affects insulin expression and antigen presentation in beta cells. When CVB5 infects the beta cell, the virus 2A protease cleaves eIF4G, leading to inhibition of cap-dependent translation of the host cell. This results in the depletion of short-lived GGA2, which delays sorting at the TGN and thereby to acidification of the Golgi complex and subsequent pre-activation of lysosomal hydrolases, such as cathepsin B and D. Cathepsin activation results in the creation of new peptides (neoantigens) in the Golgi lumen, while the lower pH affects the binding of peptides to HLA-I. Hence, neoantigens with a lower pI could displace peptides preloaded in the ER at pH 7 to in-transit HLA-I molecules, thus altering the repertoire of self-antigens presented by CVB5-infected β-cells.

The extent to which the individual different HLA-I alleles expressed in ECN90 cells contributed to presentation of the detected peptides varied between samples from infected and control cells (Suppl. Fig. 7d). In control cells, most peptides (69%) were predicted to be presented by the HLA-A alleles, namely the A*02:01 and A*03:01, contributing to 30% and 39%, respectively. This was in stark contrast to the samples from infected cells, where HLA-A alleles were predicted to cumulatively present only 38% of the peptides (A*02:01 18%; A*03:01 20%). In infected cells this was compensated by the share of peptides predicted to bind HLA-B alleles, which cumulatively accounted for 52% (compared to only 22% in control cells), with B*40:01 contributing 18% (6% in control cells) and B*49:01 presenting 34% of peptides (16% in control cells). Conversely, the estimated contribution of HLA-C alleles was similar in both conditions. Of all the alleles, B*40:01 was the one with the biggest change in the contribution of presented peptides between the two conditions. Specifically, of the 54 peptides consistently recovered from infected cells only, 25 were predicted to bind to B*49:01 (46%) and 18 to B*40:01 (33%) (Suppl. Fig. 7f). The alleles A*02:01 and A*03:01, the HLA-I most investigated in the context of T1D, were predicted to present only 4 (7%) and 3 (6%) of these 54 peptides. Strikingly, the mean isoelectric point (pI) of the unique peptides isolated from all replicates of EV-9 DM-infected ECN 90 cells was 5.20, with the mean pI of those from B*40:01 being 4,22 (Fig. 8e), compared to the mean pI of 7.36 of the unique HLA-I peptides eluted from uninfected cells, with none of them binding to HLA-B. Among these was a peptide from Chromogranin A, a SG cargo and autoantigen of T1D, with a pI of 3.79. As luminal pH affects the binding of peptides presented by HLA molecules, the presentation of immunopeptides with a lower pI in EV-9 DM-infected cells was likely skewed by the more acidic environment encountered by HLA-I in transit through the Golgi complex. As previously reported (Vecchio et al. 2025) only two of the 165 viral peptides predicted to strongly bind any of the six HLA-I alleles expressed by ECN90 cells were eluted from infected cells (Suppl. Fig. 7g). One peptide with a pI of 3.79 was conceivably presented by HLA-B B*40:01, while the other with a pI by 10 A*03:01.

## Discussion

We previously reported that CVB5 inhibits cap-dependent translation in infected insulinoma cells and mouse islets while promoting PTBP1-mediated cap-independent translation of its own uncapped RNA genome and insulin SG cargoes (Knoch et al., 2014). Despite this, in CVB5-infected cells insulin SG stores were profoundly depleted. However, the molecular mechanisms underlying these changes and their potential impact on antigen presentation by infected beta cells remained unclear.

In this study we demonstrate that similarly to CVB5 infection, expression of CVB5 protease 2A, but not 3C, is sufficient to reduce SG stores in MIN6 cells. As in other cell types, CVB5 protease 2A cleaves eIF4G, a key factor required for cap-dependent translation of host cell mRNAs (Sonenberg, 1990). We ruled out proteasomal degradation being responsible for the selective loss of mature SG cargoes upon CVB5 2A^pro+^ expression and identified the Golgi complex—specifically, the trans-Golgi network (TGN) as the secretory compartment where missorting of newly synthesized SG cargoes may lead to their premature degradation. Proteomic analyses revealed that CVB5 2A^pro^-induced blockade of cap-dependent translation rapidly depletes the amounts of short-lived GGA2, an adaptor protein essential for sorting secretory proteins at the TGN to post-Golgi compartments by binding to sorting receptors of the VSP10 family, like SorcS1 and mannose-6-phosphate receptors (M6PRs). Sorcs-1 has been implicated in the biogenesis of insulin SGs (Kebede et al. 2014), whereas M6PRs are responsible for the delivery of pro-cathepsins to lysosomes (Doray et al., 2020). Accordingly, we show that in the absence of GGA2 newly-synthesized insulin and cathepsin B, D and L are retained in the TGN. On the other hand, the levels and surface trafficking of MHC class I molecules were unaffected. Remarkably, we confirmed that in patients who succumbed shortly after the onset of T1D VP1^+^ (EV-infected) beta cells exhibited reduced levels of both GGA2 and insulin. As expected, TGN retention of V-ATPase, which is also sorted to insulin SGs and lysosomes, led to acidification of the Golgi complex lumen in both *Gga2*^*-/-*^ and CVB5 2A^pro+^ MIN6 cells, with premature activation of cathepsins B and D (Fig. 8f).

As in MIN6 cells, GGA2 was also depleted in EV9-DM-infected ECN90-beta like cells. In this instance we observed a profound alteration of the immunopeptidome presented by HLA-I. These findings are consistent with those previously reported for CVB1 or CVB3 infected ECN90-beta like cells (Vecchio et al, 2025). Also in that case HLA-B alleles contributed preferentially to the presentation of novel antigens in infected cells, with a minor representation of peptides of viral origin. Notably, we found that peptides bound to HLA-B of infected cells had a very low pI compared to those presented in HLA-A alleles, which were exclusively involved in the presentation of peptides unique to non-infected cells. Recently it has been shown that the proinflammatory cytokine interferon alpha promotes the generation and presentation of neoantigens by HLA-B alleles in human beta cells (Carre’ et al, 2025). In the future it would be interesting to test whether exposure of beta cells to interferon alpha alters pH homeostasis in their secretory pathway.

Like those originating from host cell proteins, viral antigenic peptides are generated in the cytosol by proteolysis in the proteasome and then imported into the ER to be mounted on HLA-I in the presence of a neutral pH. As HLA-I traffic through the secretory pathway, weak HLA-I binders, including viral peptides, may be lost due to the progressive acidification of the luminal pH (Saikira et al, 2022). Our findings indicate that this process may be exacerbated in the case of EV-infected beta cells due to the even greater acidification of the Golgi lumen and the generation of novel peptides by prematurely activated cathepsins. Several of these neoantigens, including some resulting from the proteolysis of insulin SG cargoes, could competitively displace HLA-I weak binders and thus be presented at the surface of the cells. As the genome of EV, such as CVB, does not encode secretory proteins, this mechanism could facilitate virus evasion from immune surveillance by replacement of viral peptides with those resulting from cathepsin-mediated cleavage of SG cargoes.

Cathepsins typically become active only once they reach the endolysosomal compartment. and in antigen presenting cells they generate peptides to be mounted on HLA-II alleles. In these cells they have also been implicated in cross-presentation, i.e., in the cleavage of endocytosed extracellular antigens for presentation to CD8^+^ T-cells by HLA-alleles. Our findings indicate that a novel sort of cross-presentation may also occur in an infected non-professional antigen presenting cells, such as beta cells, with peptides resulting from cathepsin cleavage of self-antigens in the TGN being mounted on passing HLA-I alleles. These neoantigens, in turn, may be recognized by CD8^+^ T-cells, which are mainly responsible for killing beta cells in T1D. In conclusion, our findings suggest that alterations in the pH homeostasis along the secretory pathway following viral infection or in response to other intrinsic and extrinsic factors modulate presentation of self-antigens and immune tolerance.

## Acknowledgements

We are indebted to Stefan Stefanovic for the immunopeptidome analysis of ECN90 cells. We thank, Ezio Bonifacio, Sebastian Springer, Monika Füssel and Peter Cresswell, as well as all members of the Solimena lab for discussion and advice; Frank Möller for creating the scheme in figure 9 and Mrs. Katja Pfriem for administrative support; the Light Microscopy Facility a Core Facility of the CMCB Technology platform at TU Dresden and the CMCB Flow Cytometry Core Facility. This work was supported by the German Center for Diabetes Research (DZD e.V.), which is financed by the German Ministry for Education and Research and from the Innovative Medicines Initiative 2 Joint Undertaking under grant agreements no. 115881 (RHAPSODY) to M.S.; by INNODIA to M.S., R.S., R.M, S.J.R. and N.M. INNODIA has received funding from the Innovative Medicines Initiative 2 Joint Undertaking under grant agreement on. 115797 (INNODIA). This Joint Undertaking receives support from the Union’s Horizon 2020 research and innovation programme, ‘EFPIA’, ‘JDRF’ and ‘The Leona M. and Harry B. Helmsley Charitable Trust’. This work was also supported by a Steve Morgan Foundation Grand Challenge Senior Research Fellowship to S.J.R (22/0006504). Views and opinions expressed are those of the authors only and don’t necessarily reflect those of the funding agencies. Neither the European Union nor any of the granting authorities can be held responsible for them.

## Author contributions

Conceptualization, K.-P.K. and M.S.; methodology, K.-P.K.; validation, K.-P.K., I.K., Y.K., J.M.-P. and M.G.; formal analysis, M.S.; K.-P.K., A.K., C.L., I.K., Y.K., A.P., C.W., A.S., K.G., A.M., M.G., J.M.-P., A.S.; resources, A.C., R.S., R.M., S.H., N.G.M., S.J.R. and M.S.; writing – original draft, K.-P.K. and M.S., review & editing, K.-P.K., R.S., S.J.R., N.G.M., M.S.; visualization, K.-P.K.; funding acquisition, M.S., R.S., R.M., S.J.R; N.G.M.

## Material and Methods

### Cell culture

Mouse insulinoma MIN6 cells were cultured as described (Ishihara et al. 1993). For Insulin secretion assay cells were kept in resting medium for hour before being stimulated for 2 hours by the addition of 25 mM glucose (Ort, et al. 2001). After 2hrs stimulation cells were harvested. Human pancreatic ECN90 cells (Rachdi et al., 2020) were kindly provided by Dr. Raphael Scharfmann (Institut Cochin, Paris, France) and cultured as described (Gonzalez-Duque et al. 2018). CVB5 strain Falkner and Echo-9 virus strain DM used in this study were kindly provided by Dr. Merja Roivainen (National Institute for Health and Welfare, Helsinki, Finland). Infection and harvesting of the cells was done as described (Knoch et al. 2014). For the measurement of cathepsin activity cells were harvested in cell lysis buffer provided in the kits (Abcam). Enzymatic activity was measured according to the manuals.

### Cloning

The cDNA of 2A and 3C protease were amplified by PCR and cloned into pcDNA3.1 (INVITROGEN) following standard protocols. The cDNA for CtsB, CtsD and CtsL as well as the Atp6v0d1 was cloned into the pEGFP-N1 vector (Clontech). For CRISPR/Cas9 the mouse Gga2 specific sequence 5’-AGGGCCGCCCTCCGCCGACT was cloned into pSpCas9(BB)-2A-GFP vector according to Ran et al. 2013. pSpCas9(BB)-2A-GFP (PX458) was a gift from Feng Zhang (Addgene plasmid # 48138 ; http://n2t.net/addgene:48138RRID:Addgene_48138). For measurement of the organelle specific pH we cloned CFP into the pEGFP vector to replaced eGFP. The organelle specific proteins were cloned after that in this CFP containing plasmid as shown in Supplemented figure 7.

### Protein analysis – western blotting

MIN6 cells or ECN90 cells were lysed as described (Ort et al. 2001). Protein concentration in the detergent soluble material was measured by Pierce™ BCA Protein Assay Kit (ThermoFisher). Cell extracts were separated by SDS-PAGE and immunoblotted as described (Knoch 2004). Antibodies used are listed in Suppl. Table 2. Chemiluminescence was performed using the Supersignal West Pico Substrate (Pierce) and detected with an Amersham IMAGER 600 System (GE Healthcare). Detected signals were quantified using ImageQuant TL software (GE Healthcare). For quantification of the maturation status metabolic labeling of MIN6 cells was done using 100 µCi ^35^S-methionine/35 mm well for 2 hours. After 5 times washing the cells were extracted and Ica512 and Pc1/3 were isolated by immunoprecipitation. Precipitated protein was loaded on a 8% SDS-PAGE and blotted on nitrocellulose. Radioactive signals were captured with a BAS 1800II phosphoimager (Fuji) using the Image Gauge v3.45 software (Knoch et al. 2004).

### Real time PCR

Total RNA from MIN6 cells was prepared according to the RNeasy Kit (QIAGEN). 1µg total RNA was reverse transcribed with SuperScript II reverse transcriptase (Invitrogen) and oligo d(T) primer. mRNA levels were measured by quantitative real-time PCR with the GoTaq qPCR Master Mix Kit (PROMEGA) and a AriaMx Thermocycler (Agilent Technologies). Primer used for this purpose are listed in Suppl. Table 3. Normalization of real-time PCR data was performed by parallel amplification of *ß-Actin* mRNA, as described in Knoch et al., 2014.

### Immunocytochemistry

MIN6 cells were fixed with 4% paraformaldehyde and permeabilized with 0.2% saponin according to Knoch et al. 2004. Fixed cells were immediately stained using the designated antibodies listed in Suppl. Table 2. Images were acquired with a LSM880 Airyscan (ZEISS) and processed with ZEN 3.0 software (ZEISS).

For multiplex tissue immunofluorescence, FFPE pancreas tissue sections were obtained from the Exeter Archival Diabetes Biobank (EADB; https://pancreatlas.org/) with ethical permission from the West of Scotland Research Ethics Committee (ref: 15/WS/0258), or the network for Pancreatic Organ Donors with Diabetes (nPOD) (Suppl. Table 1). Sections were dewaxed in histoclear and rehydrated in degrading ethanols (100%, 90%, 70%, methanol. Heat-induced epitope retrieval (HIER; 10mM citrate pH6.0) was performed to unmask the epitopes by placing the sections in a pressure cooker in a microwave oven on full power for 20 mins. The sections were then blocked with 5% normal goat serum (NGS) and sequentially incubated with the relevant primary antibodies (Suppl. Table 2), before being probed with an appropriate Alexa Fluor secondary antibody (Suppl. Table 2). The sections were counterstained with DAPI and mounted for fluorescent microscopy performed using the Leica DMi8 SP8 LIGHTNING modular (Leica Microsystems, Milton Keynes, UK) confocal microscope. Quantification was performed using the Indica HALO image analysis platform. The area quantification module was used to measure the mean fluorescent intensity (MFI) of manually annotated regions of interest (ROIs) (2.9µm^2^). The ROIs were randomly selected within VP1 positive insulin-containing cells or VP1 negative insulin-containing beta cells. The summary data were exported in Excel format and analysed in RStudio.

For Live cell imaging MIN-6 cells were transiently transfected by Insulin-SNAP plasmid and stained after 30min SNAP-Cell Block treatment with SNAP-Cell-TMR substrate as described in [Ivanova et al. 2012]. After labeling cells were washed three times for 30min with medium. Labeling procedures were performed at 19°C, when insulin granules exit from Golgi is blocked insulin. Then, cells were placed under microscope in Warner heating chamber with stabilized 37°C and CO_2_ supply for live imaging (Andor Spinning Disc based on Andor Eclipse Ti inverted stand confocal microscope and Yokogawa, CSU-X1 spinning disk equipped by a 100x/1.45 Plan Apochromat oil immersion objective Nikon, fast piezo z-drive stage and Andor iXon EM+ DU-897 BV back illuminated EMCCD camera). 3-4 viewfields with cells were selected. The microscope was programmed to visit a selected viewfield every 25 seconds and acquire 5 time-sequential z-stacks (11 planes, z-step 0.5µm) with a fast frame rate (12 planes per second). The images were deconvolved and individual granules were segmented by MotionTracking software (http://motiontracking.mpi-cbg.de). In total, 21 movies (64 cells) in control and 24 movies (83 cells) in GGA2 KO conditions were recorded and analyzed. The mean peer-to-peer distance between every couple of granules was calculated. The course of mean distances grew when granules spread in the cell and later stabilised. The mean peer-to-peer distance growth is well described by two straight lines: one describes the granule spread and second, horizontal, the final granule distribution. Therefore we used a Bayesian approach to fit two lines with free-to-find transition time points. The fitting procedure is implemented in MotionTracking software. The size of granule was calculated as a diameter of the projection of 3D segmented granule on X-Y plane.

For Fluorescence Lifetime Imaging Microscopy a SP8 Falcon microscope was used. Following the protocol from Neukamp (2017) the measurement of the CFP half live transfected cells were kept in KCl-rich media pH 7,4 as descripted (Kim et al. 1996). For the standard curve MIN6 cells were incubated for 5min in KCl-rich buffer with a pH range of 5,5-7,5 plus 5µM Nigericin and 5µM Monensin.

### Electron microscopy

MIN6 wt and Gga2^-/-^ cells cultured in 6 well plates were fixed for electron microscopy processing with 4% PFA and 2.5 % glutaraldehyde in sodium phosphate buffer. The samples were further processed according to a protocol modified from (Seligman et al. 1966) using osmium tetroxide (OsO_4_), thiocarbohydrazide (TCH) and OsO_4_. In brief, samples were incubated in a 2% aqueous OsO_4_ solution containing 1.5% potassium ferrocyanide and 2 mmoll^−1^ CaCl_2_ (30 min on ice), washed in water, 1% TCH in water, rinsed in water and incubated again in 2% OsO_4_. After washing the samples were en-bloc stained with 1% uranyl acetate (UA), washed in water and dehydrated in a graded series of ethanol. The samples were embedded into epoxy resin Embed812 (Science Services). After curing of the resin at 60°C sections with a thickness of 70 nm were cut. Poststaining of the sections was done with 2% UA and 1% lead citrate in H_2_O. The sections were imaged in a 80kV JEM1400Plus transmission electron microscope (JEOL).

### FACS sorting

MIN6 WT and *Gga2*^-/-^ cells were pre-incubated with 125 U/ml IFN-γ in MIN6 medium for 72 hours to induce MHC class I expression prior to FACS analysis. Different temperature conditions were used to modulate protein export from the Golgi complex: constant incubation at 37°C, 1.5 hours at 19°C, and a combination of 1.5 hours at 19°C followed by 2 hours at 37°C. Cell detachment was achieved using 20 mM EDTA in DPBS, followed by a 5-minute incubation at either 37°C or room temperature. After washing cells were resuspended in FACS buffer containing 1 mM EDTA, 25 mM HEPES pH 7.0, 0.1% Albumin in PBS.

After staining for MHC classI single cell fluorescence was measured using an LSRFortessa™ Cell Analyzer (BD Biosciences) with FACSDiva Software (BD Biosciences). 10,000 events were recorded for each experimental condition. Data analysis was performed using FlowJo™ software. Fluorescence intensity of the H2kb signal was assessed in FACS experiments conducted with six biological replicates. Statistical analysis was performed using R software and the lme4 package, using the lmer function to construct a linear mixed-effects model.

### RNA interference

Short-interfering double stranded RNA (siRNA) oligonucleotides for Gga2 and Rab3b were synthesized with the Silencer siRNA Construction Kit (Ambion). Primers are reported in Suppl. Table 3. Control scrambled siRNA oligonucleotides were previously described (Knoch et al. 2006).

### LC-MS/MS and label-free quantitative analysis

Ten micrograms of MIN6 cell lysates were digested using a modified FASP procedure (Wisniewski et al. 2009). Briefly, proteins were reduced and alkylated by DTT and iodoacetamide prior to dilution with urea to a final concentration of 4M. Samples were then centrifuged through a 30kDa cut-off filter (Sartorius) and washed twice with 8 M urea in 0.1M Tris/HCl pH 8.5 and 50mM (NH_4_)HCO_3_. Proteolysis was performed on the filter using Lyc-C (Wako Chemicals) and trypsin (Promega). Resulting peptides were eluted and acidified with trifluoroacetic acid. LC-MSMS analysis was performed on a QExactive HF mass spectrometer (ThermoFisher Scientific) online coupled to a UItimate 3000 RSLC nano-HPLC (Dionex). Acquired raw data were loaded to the Progenesis QI software (Nonlinear Dynamics, Waters). After alignment, filtering and normalization, all MSMS spectra were exported and searched against the Swissprot mouse database (16871 sequences, Release 2016_02) using the Mascot search engine. A Mascot-integrated decoy database search calculated an average false discovery of < 1% when searches were performed with a percolator algorithm and a significance threshold of 0.05. Peptide assignments were re-imported into the Progenesis QI software. The abundances of all unique peptides allocated to each protein were summed up. Resulting normalized abundances of the single proteins in the individual samples were then used for calculation of fold-changes of proteins and significance values by a Student’s T-test.

### Genomic DNA isolation and HLA genetic haplotyping

Genomic DNA of ECN90 cells was isolated for HLA haplotyping using the DNeasy Blood and Tissue kit (Qiagen) according to the manufacturer’s instructions. HLA genotyping was done by the DKMS Life Science Lab GmbH, Dresden, Germany.

### HLA ligand analysis

The HLA class I of the human beta cells were isolated using standard immunoaffinity purification (Falk et al., 1991). HLA–peptide complexes were eluted by repeated addition of 0.2% trifluoroacetic acid (Merck). Eluted HLA ligands were purified by ultrafiltration using Amicon Centrifugal Filter Units (Millipore). Isolated peptides were separated by reversed-phase liquid chromatography (nano-UHPLC, UltiMate 3000 RSLCnano; Thermo Fisher Scientific) and analyzed in an online-coupled Orbitrap Fusion Lumos mass spectrometer (Thermo Fisher Scientific). Samples were analyzed in five technical replicates and sample shares of 20% trapped on a 75 µm Å∼ 2 cm trapping column Acclaim PepMap RSLC (Thermo Fisher Scientific). For HLA class the mass range was limited to 400–650 m/z. LC-MS/MS results were processed using Proteome Discoverer (v.1.3; Thermo Fisher Scientific) to perform database search using the Sequest search engine (Thermo Fisher Scientific) and viral EV-9 DM and the human proteome as reference database annotated by the UniProtKB/Swiss-Prot (http://www.uniprot.org), status September 27th 2013 containing 20,279 reviewed sequences. The isoelectric point of the isolated peptides were calculated using Compute pI/Mw tool at Expasy.org

### Statistics

GraphPad Prism version 10 was used for statistical analyses. p values were calculated with the one-way ANOVA t-test and indicated in the figures as follows: *p<0.05, **p<0.01 and ***p<0.001. Error bars show standard deviations from at least three independent experiments. For the graphical representation GraphPad Prism and RStudio ggblot2 were used.

## Supplementary Material

### Supplementary Figure and Table Legends

**Suppl. Figure 1:**
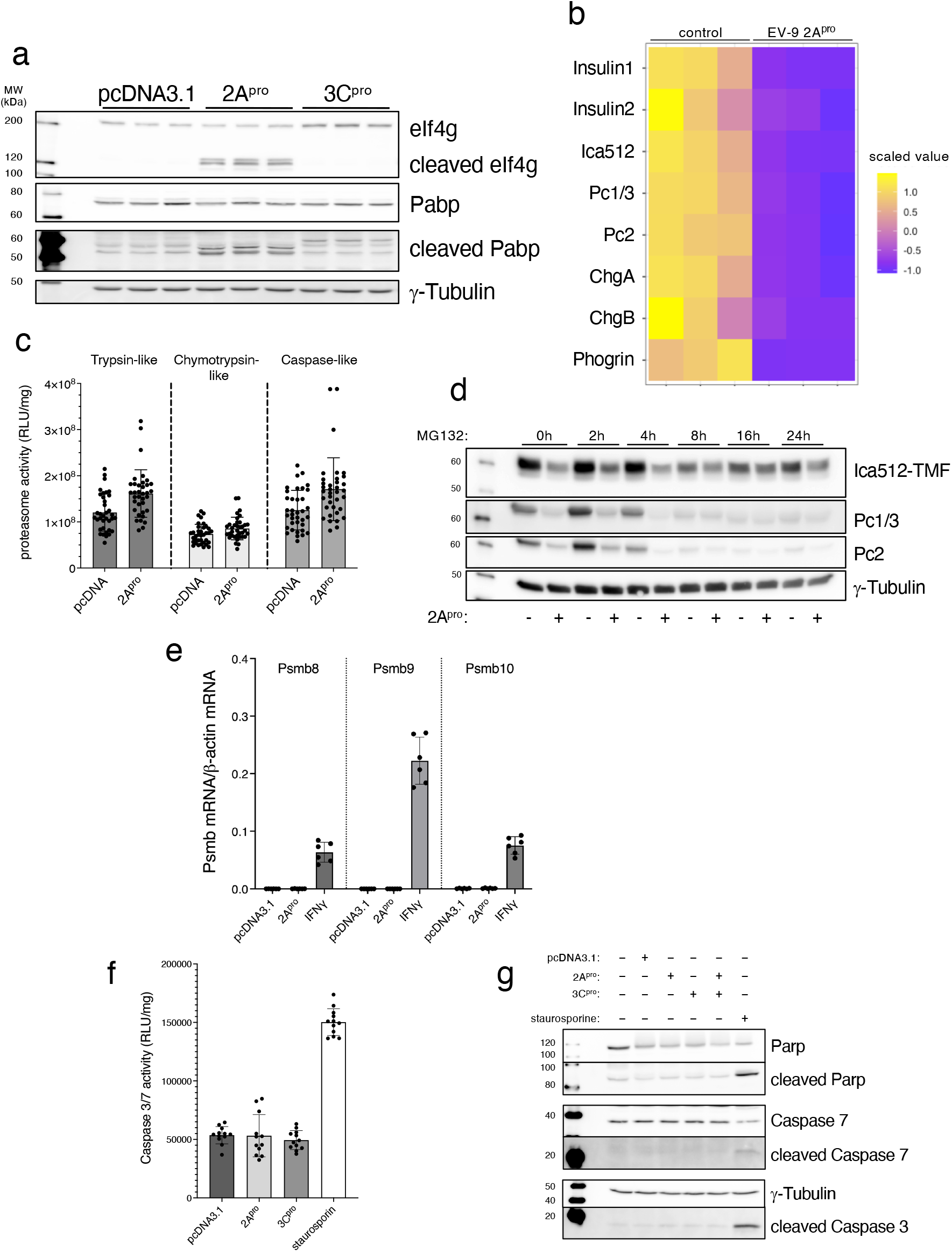
CVB5 2A, but not 3C, depletes insulin granule stores, but does not change proteasome activity nor induces apoptosis. (a) Western blot for eIf4G, Pabp, and γ-Tubulin in extracts of MIN6 cells transfected with either CVB5 2A or/and 3C proteases or the empty vector pcDNA3.1. (b) Heatmap for Insulin1, Insulin2, Ica512, Pc1/3, Pc2, Chga and Chgb and Ptpn2 from global proteomic analysis (Z-score clustered) of extracts of MIN6 cells transfected with EV-9 DM 2A^pro^ or pcDNA3.1 alone. Data are from 3 independent experiments. (c) Quantification of the trypsin-like, the chymotrypsin-like and caspase-like proteasome activity in extracts of MIN6 cells transfected with either CVB5 2A^pro^ or pcDNA3.1 alone. (d) Western blot for Ica512, Pc1/3, PC2 and γ-Tubulin in MG131 treated extracts of MIN6 cells transfected with either CVB5 2A^pro^ or pcDNA3.1 alone. (e) Quantitative RT-PCR for *Psmb8, Psmb9* and *Psmb10* in MIN6 cells transfected with CVB 2A^pro^ or pcDNA3.1 alone, as well as in IFNγ treated MIN6 cells. mRNA expression levels were normalized by β-actin mRNA. (f) Caspase 3/7 activity in MIN6 cell extracts overexpressing CVB 2A^pro^ or 3C^pro^ or staurosporin treated untransfected cells. (g) Western blot for Parp, Caspase7, Caspase3 and γ-Tubulin in extracts of MIN6 cells transfected with either CVB5 2A^pro^ 3C^pro^ or pcDNA3.1 alone. Staurosporin treated MIN6 cell extracts were used as positive control for caspase cleavage in cells undergoing apoptosis All data are from ≥ 3 independent experiments

**Suppl. Figure 2:**
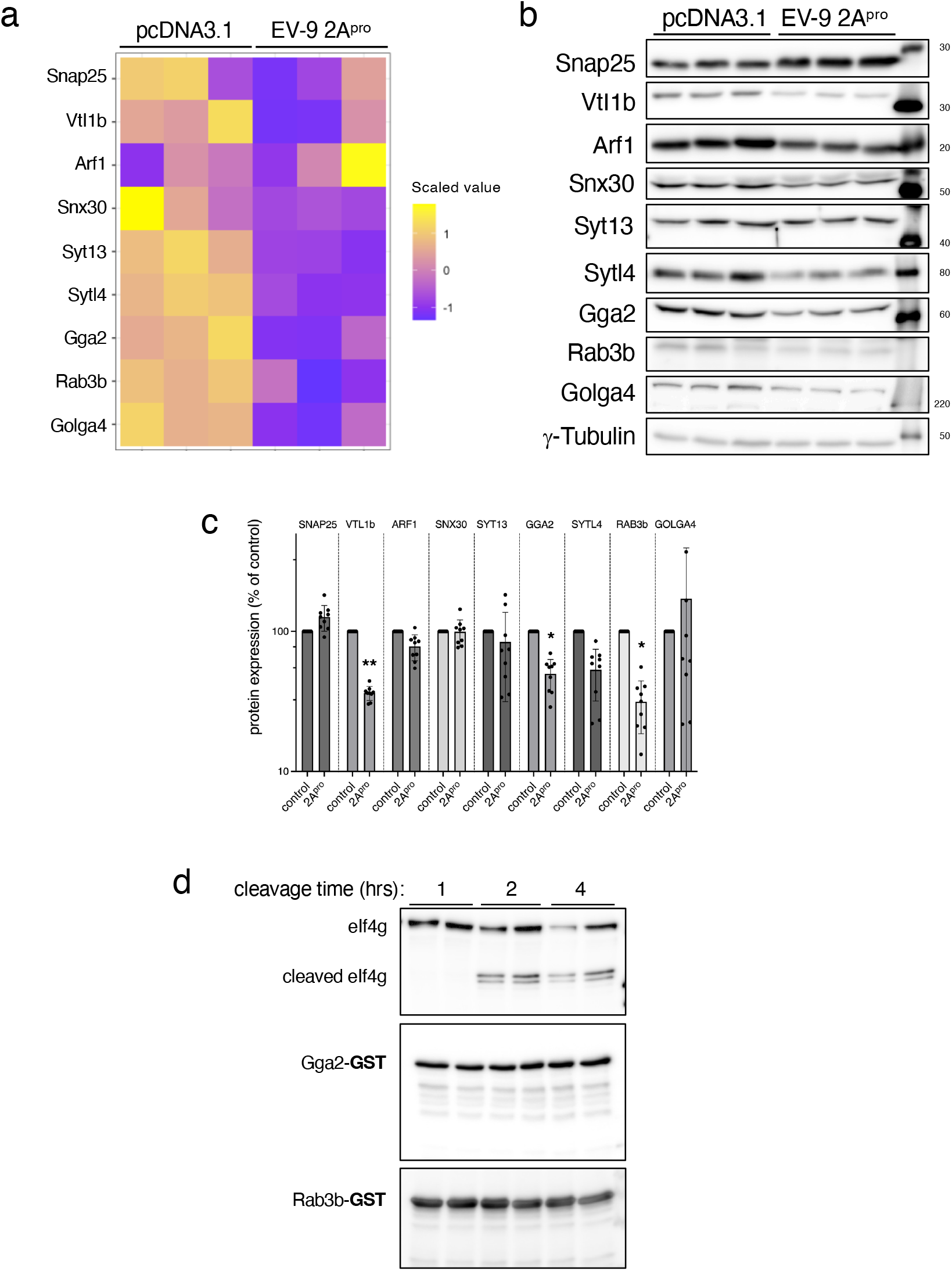
Components of the post-Golgi trafficking machinery are not direct substrates of 2A^pro^. (a) Heatmap for selected players in post-Golgi membrane trafficking as revealed by global proteomic analysis (Z-score clustered) of extracts of MIN6 cells transfected with EV-9 DM 2A^pro^ or pcDNA3.1 alone. (b, c) Western blots (b) and corresponding quantifications (c) for selected post-Golgi trafficking proteins downregulated upon expression of EV-9 DM 2A^pro^. Data are from ≥ 3 independent experiments; statistical values were calculated by one-way ANOVA *p<0.05. (d) Western blots for eIf4g and GST of CVB5 2A^pro^ digest reaction. Cell extracts or purified Gga2-GST or Rab3b-GST were used as controls.

**Suppl. Figure 3:**
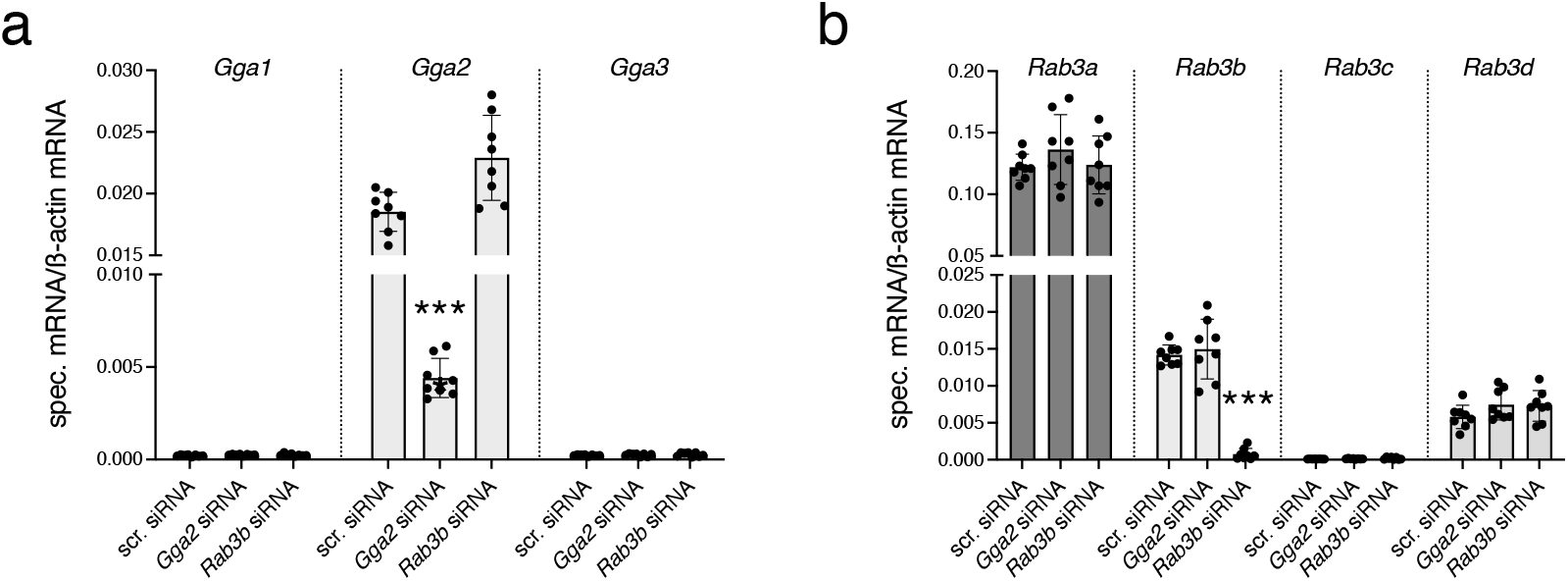
*Gga2* reduction did not lead to a change in expression of Gga1 and Gga3. Quantitative RT-PCR for (a) *Gga1, Gga2* and *Gga3* and (b) *Rab3a, Rab3b Rab3c* and *Rab3d* in MIN6 cells 3 days after transfection of siRNAs for *Gga2 Rab3b* or a scrambled control oligonucleotide. mRNA expression levels were normalised by β-actin mRNA. All data are from ≥ 3 independent experiments

**Suppl. Figure 4:**
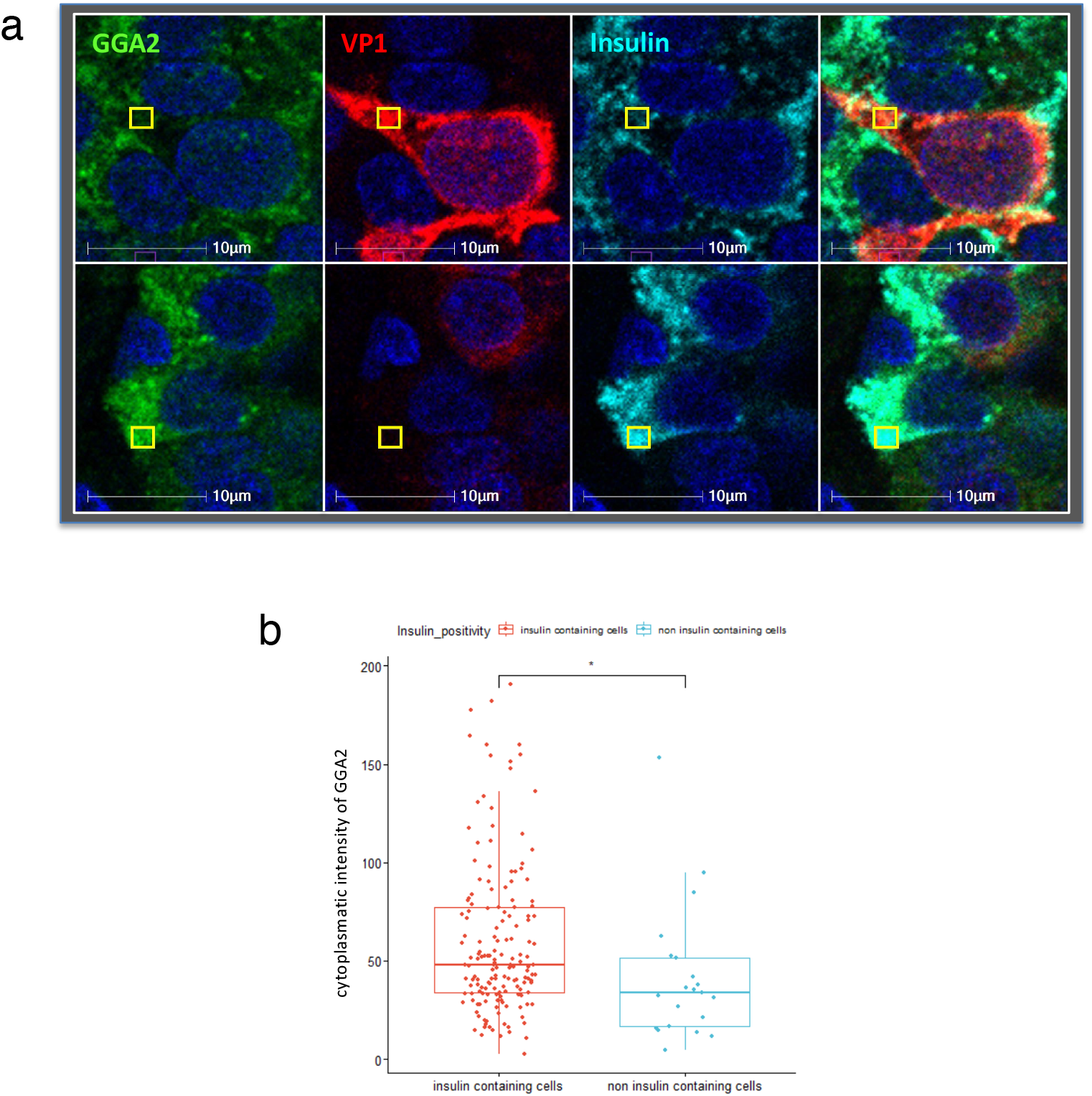
GGA2 expression is increased in insulin positive cells in human pancreatic tissue. (a) Immunostaining of insulin (cyan), GGA2 (green) and VP1 (red) in human tissue sections. Confocal images were inputted into HALO. Using an annotation tool, uniform square-shaped annotations were created. These were randomly annotated in 8-9 areas in VP1^+^ beta cells (representative image in bottom panel). Using the area quantification module within HALO, the average GGA2 staining intensity was calculated and presented in the box plot. (b) Quantification of GGA2 in insulin containing or insulin negative cells.

**Suppl. Figure 5:**
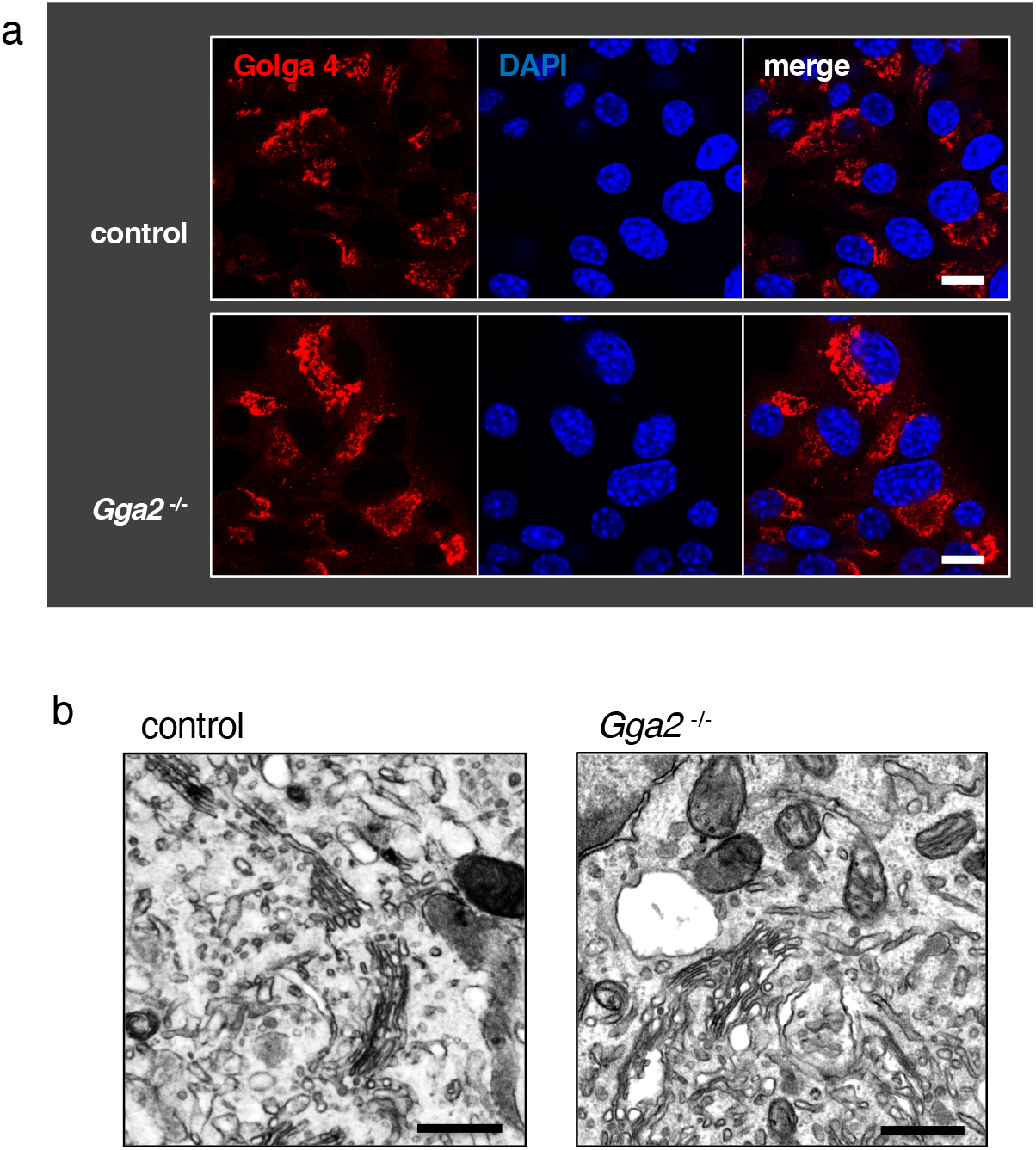
The Golgi apparatus shows no morphological change upon knockout of *Gga2*. (a) Immunostaining of Golga4 in wild-type and *Gga2*^-/-^ MIN6 cells. Nuclei were counterstained with DAPI. Scale bars = 10 µm (b) Electron microscopy picture of wild-type and *Gga2*^-/-^ MIN6 cells. Scale bars = 1µm.

**Suppl. Figure 6:**
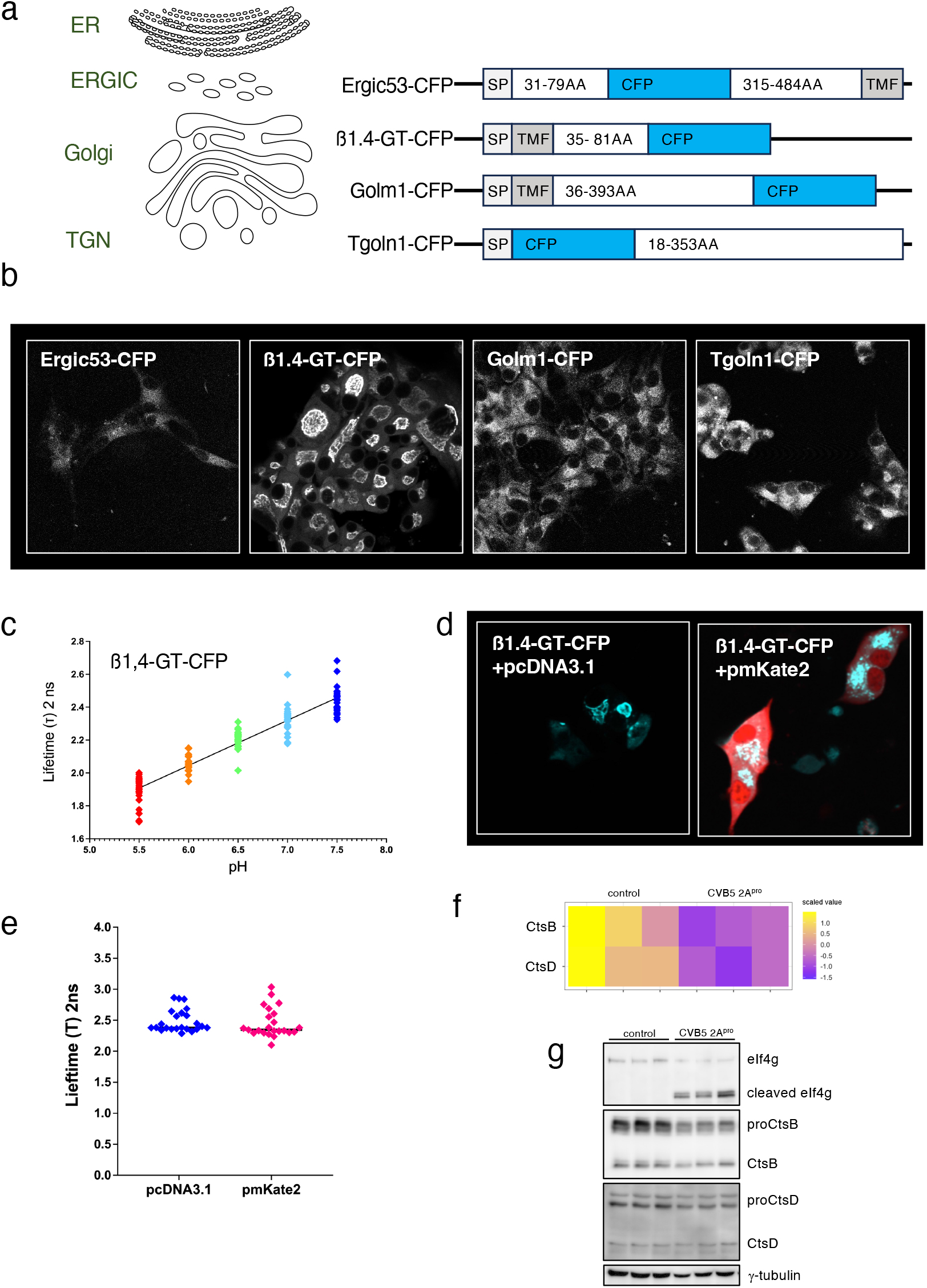
pH measurement of different cell compartments in MIN6 cells. (a) Scheme of the different CFP-containing constructs used to measure the pH in the ERGIC, cis-Golgi, late Golgi and TGN of stable transfected MIN6 cells. (b) Confocal microscopy for Ergic53-CFP, β1,4-GT-CFP, Golm1-CFP and Tgoln1-CFP stable expressed in MIN6 cells. (c) Standard curve for a pH depended half-life measurement of CFP in the range of a pH of 5.5 to 7.5 (d) Confocal microscopy for β1,4-GT-CFP (cyan) co-expressed with pmKate2 (red) in MIN6 cells. (e) Measurement of the fluorescence life-time of β1,4-GT-CFP with or without pmKate2. Data are from ≥ 5 independent experiments. (f) Heatmap for CtsB and CtsD from global proteomic analysis (Z-score clustered) of extracts of MIN6 cells transfected with CVB 2A^pro^ or pcDNA3.1 alone. (g) Western blot for eIf4G, CtsB, CtsD, and γ-Tubulin in extracts of MIN6 cells transfected with either CVB5 2A^pro^ or the empty vector pcDNA3.1.

**Suppl. Figure 7:**
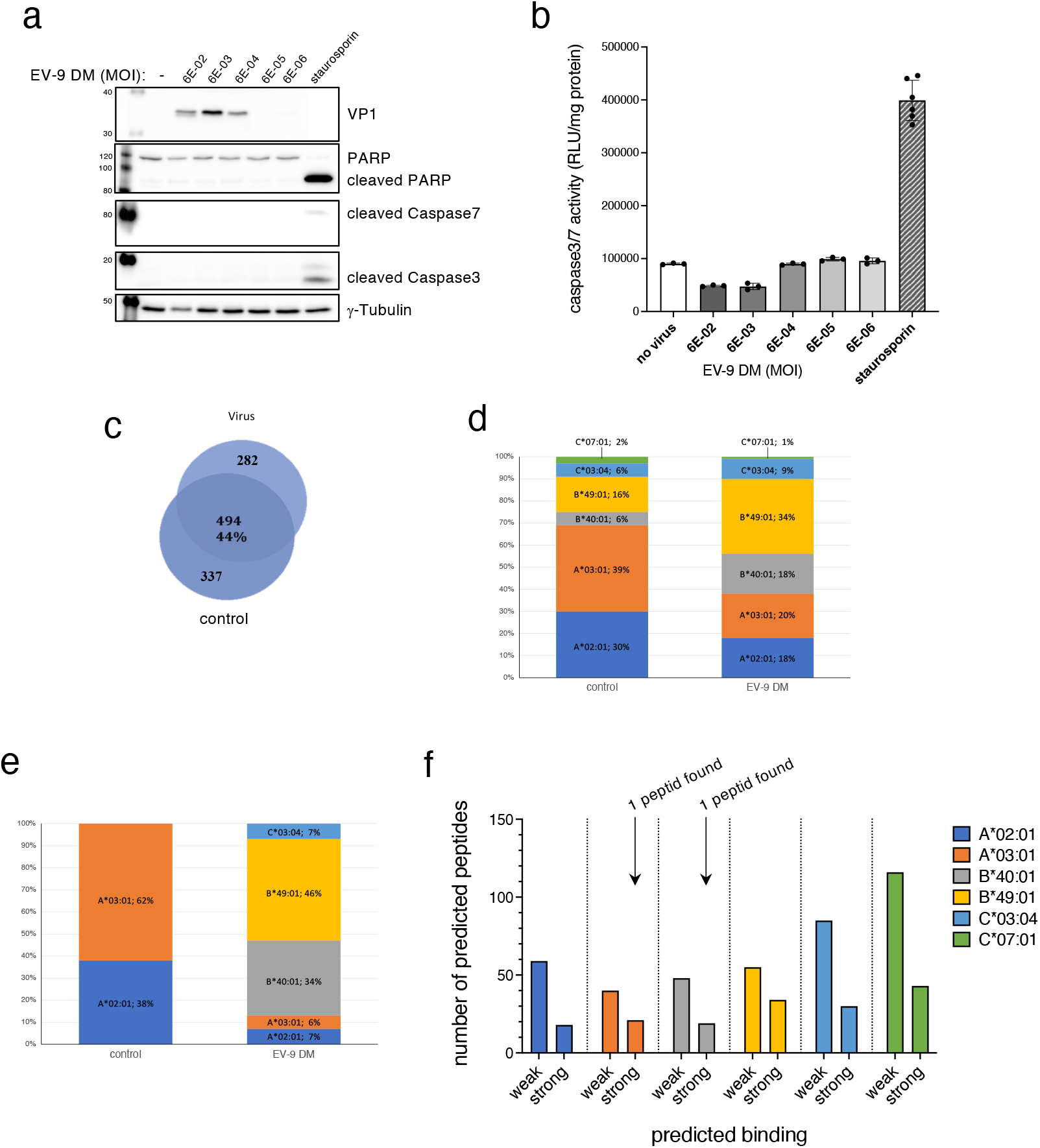
EV-9 DM infection does not induce apoptosis in human ECN90 insulinoma cells. ECN90 cells were infected with EV-9 DM virus and harvested 2 days after infection. (a) Western blot for VP1, PARP, Caspase 7, Caspase 3 and γ-Tubulin in extracts of ECN90 cells infected with different MOI of EV-9 DM. (b) Caspase 3/7 activity in ECN90 cell extracts infected with different MOI of EV-9 DM or in extract from uninfected or in staurosporin treated cells. Data are from 3 independent experiments, statistical values were calculated by one-way ANOVA. *p<0.05. (c) Venn diagrams of the number of identified HLA-I presented peptides across the two investigated conditions, EV-infected and control ECN90 cells. (d, e) Contribution of individual HLA-I alleles expressed in ECN90 cells to presentation of identified peptides. Percentage shows the ratio of peptides presented by the specific HLA-I allele relative to the total number of detected peptides in all replicates of one condition (infected ECN90 cells or control) (d) as well the ratio of peptides presented by the specific HLA-I allele relative to the number of detected peptides in all replicates of one condition (infected ECN90 cells or control) (e). (f) Numbers of predicted EV-9 DM peptides binding to HLA of ECN90 cells. Prediction was done using NetMHCpan version 4.0. (https://services.healthtech.dtu.dk/services/NetMHCpan-4.0/)

**Suppl. Table 1:** Pancreas Donor Details.

| **Donor group** | **nPOD/EADB case RRID** | **Sex** | **Age (yrs)** | **T1D Duration (yrs)** |
| --- | --- | --- | --- | --- |
| T1D | EADB E560 | F | 42 | 01. Mai |
| T1D | EADB E547 | F | 7 | At onset |
| T1D | EADB E556A | M | 18 | 0.333 |
| T1D | nPOD 6245<br>SAMN15879301 | M | 22 | 7 |
| ND | EADB 65/71 | F | 42 | N/A |
| ND | EADB 315/89 | M | 9 | N/A |
| ND | EADB 333/66 | M | 16 | N/A |
| ND | nPOD 6160<br>SAMN15879216 | M | 22. Jan | N/A |

**Suppl. Table 2:**
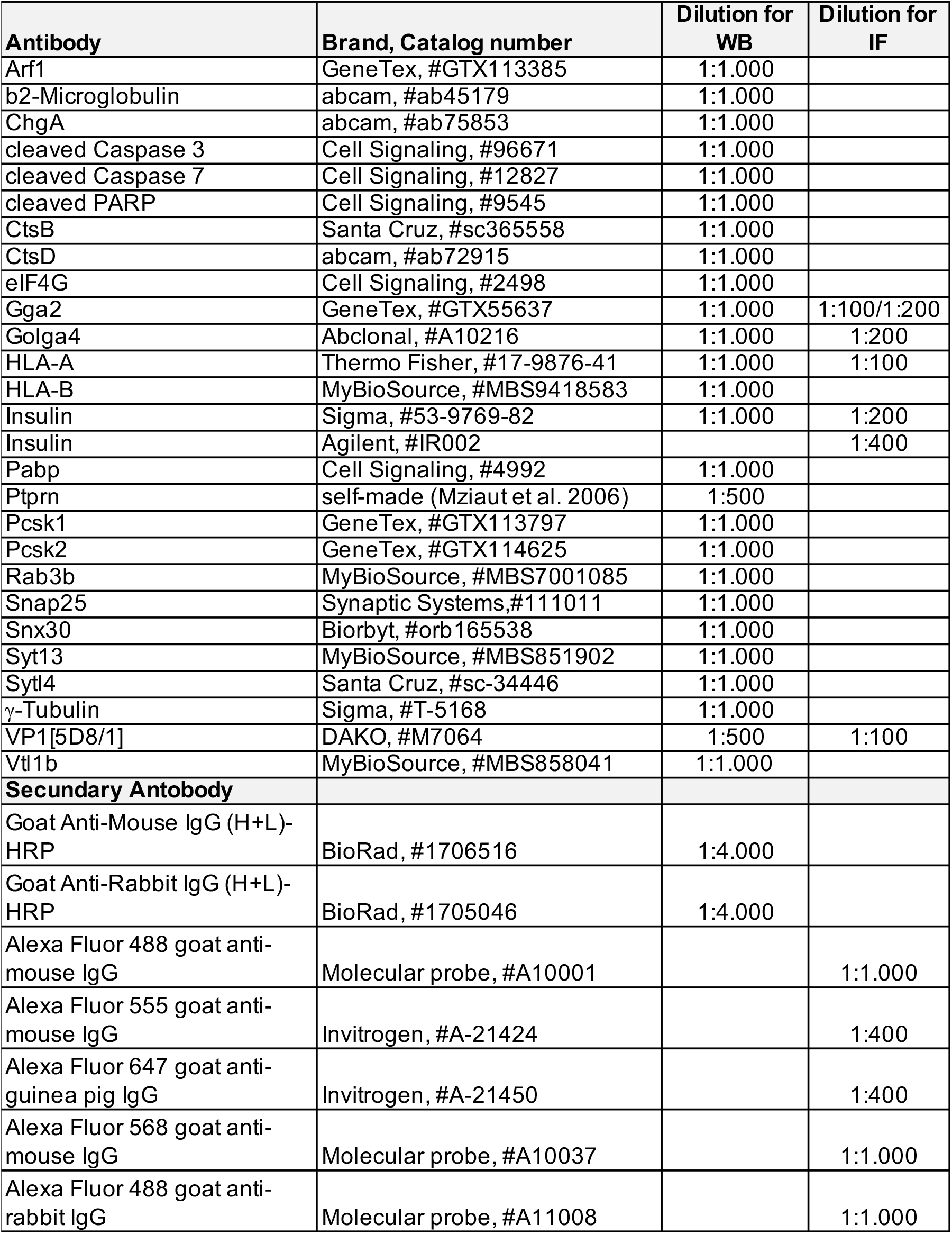
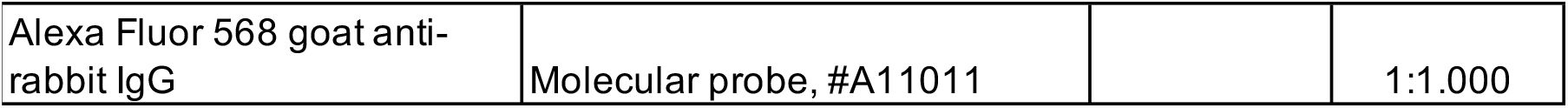
List of antibodies.

**Suppl. Table 3:**
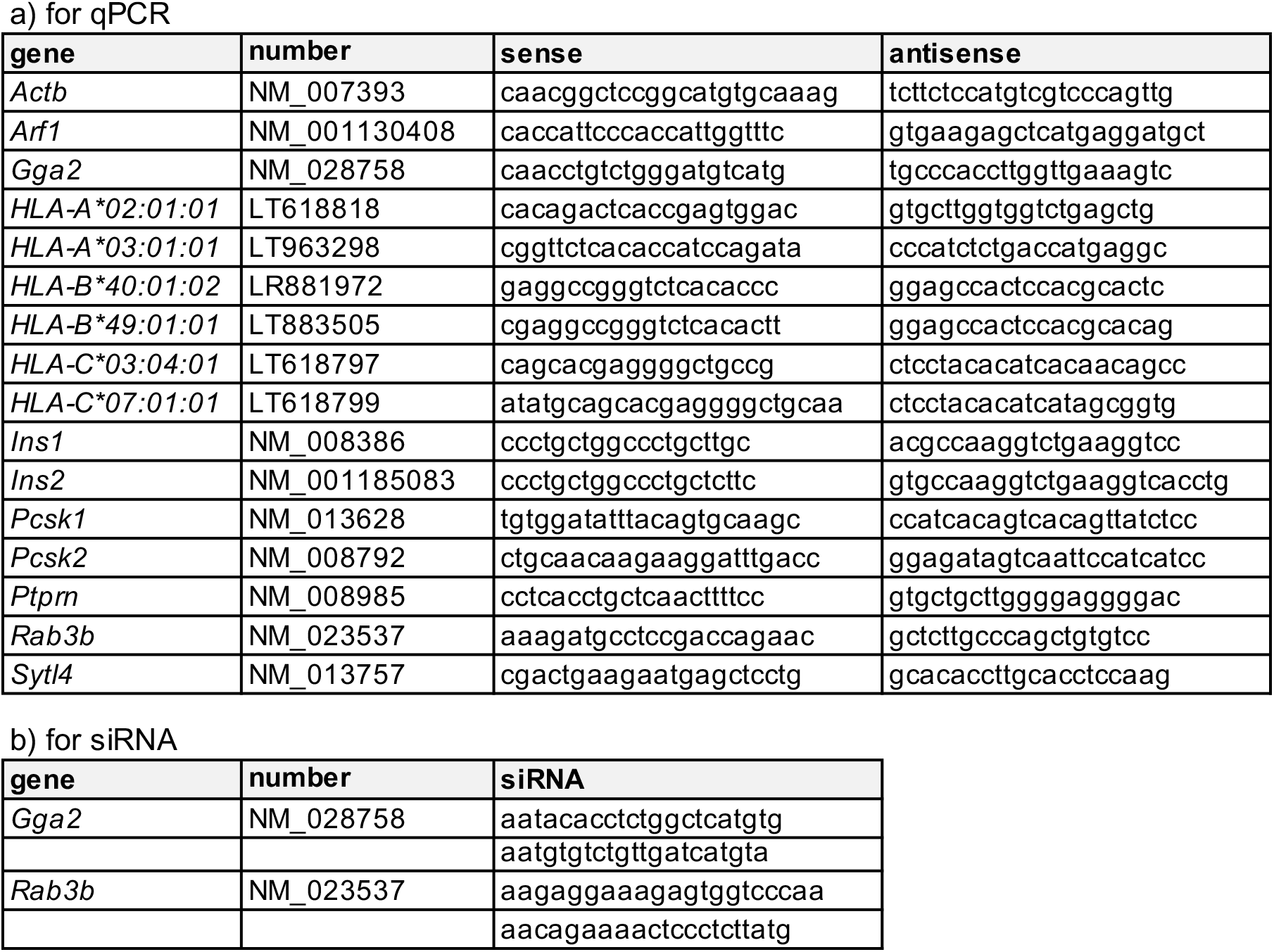
List of oligonucleotides.

## Notes

### Competing Interest Statement

The authors have declared no competing interest.

